# Designing and Comparing Optimised Pseudo-Continuous Arterial Spin Labelling Protocols for Measurement of Cerebral Blood Flow

**DOI:** 10.1101/2020.03.02.973537

**Authors:** Joseph G. Woods, Michael A. Chappell, Thomas W. Okell

**Affiliations:** Wellcome Centre for Integrative Neuroimaging, FMRIB, Nuffield Department of Clinical Neurosciences, University of Oxford, Oxford, United Kingdom; Department of Radiology, University of California, San Diego, La Jolla, CA, USA; Institute of Biomedical Engineering, Department of Engineering, University of Oxford, Oxford, United Kingdom

## Abstract

Arterial Spin Labelling (ASL) is a non-invasive, non-contrast, perfusion imaging technique which is inherently SNR limited. It is, therefore, important to carefully design scan protocols to ensure accurate measurements. Many pseudo-continuous ASL (PCASL) protocol designs have been proposed for measuring cerebral blood flow (CBF), but it has not yet been demonstrated which design offers the most accurate and repeatable CBF measurements. In this work, a wide range of literature PCASL protocols, including single-delay, sequential and time-encoded multi-timepoint protocols, and several novel protocol designs, which are hybrids of time-encoded and sequential multi-timepoint protocols, were first optimised using a Cramér-Rao Lower Bound framework and then compared for CBF accuracy and repeatability using Monte Carlo simulations and in vivo experiments. It was found that several multi-timepoint protocols produced more confident, accurate, and repeatable CBF estimates than the single-delay protocol, while also generating maps of arterial transit time. One of the novel hybrid protocols, Hybrid*_T1_*_-adj_, was found to produce the most confident, accurate and repeatable CBF estimates of all protocols tested in both simulations and in vivo (24%, 47%, and 28% more confident, accurate, and repeatable than single-PLD in vivo). The Hybrid*_T1_*_-adj_ protocol makes use of the best aspects of both time-encoded and sequential multi-timepoint protocols and should be a useful tool for accurately and efficiently measuring CBF.

## 2. Introduction

Arterial spin labelling (ASL) MRI employs magnetically labelled arterial blood as an endogenous tracer which can be used to map cerebral blood flow (CBF) (Detre et al., 1992; Williams et al., 1992). The longitudinal magnetisation of upstream arterial blood is typically labelled by inversion and, after a delay for tracer inflow (Alsop and Detre, 1996), is imaged. A single image, or multiple images using different delays, can be acquired and, with the use of a control image and an appropriate signal model (Buxton et al., 1998), the local CBF can be estimated.

A consensus paper from the ISMRM Perfusion Study Group and the European ASL in Dementia consortium recommended using pseudo-continuous ASL (PCASL) labelling with a single-PLD (post labelling delay) protocol for clinical applications, due to the superior SNR of PCASL labelling and the robustness and simplicity of using a single-PLD (Alsop et al., 2015). The PLD must be set long enough to ensure complete arrival of the labelled blood across the whole brain, while being kept short enough to preserve SNR. This leads to brain regions with short arterial transit times (ATTs) having a sub-optimally long PLD, while any regions with unexpectedly long ATTs incorrectly appear hypoperfused.

Sequential multi-PLD (Alsop and Detre, 1996) and multi-LD/PLD (label duration) (Borogovac et al., 2010; Johnston et al., 2015; Zhao et al., 2015) protocols can be used to sample the dynamics of the tracer signal, providing greater robustness of CBF estimates to variations in ATT across brain regions and subjects as well as generating potentially useful ATT maps (MacIntosh et al., 2012). However, it is often assumed that the reduction in data averaging when using multi-PLD protocols (required when acquiring multi-PLD data in a matched scan time with a single-PLD protocol) leads to a reduction in the precision of the multi-PLD CBF estimates (Alsop et al., 2015; Dai et al., 2017; Günther, 2007; Teeuwisse et al., 2014), which could outweigh the benefits of correcting for ATT effects.

We recently demonstrated that a sequential multi-PLD PCASL protocol can be objectively optimised to maintain higher CBF accuracy across a wider range of ATTs than a single-PLD or evenly spaced multi-PLD protocol (Woods et al., 2019). This is due to an improved balance between early sampling of the tracer kinetics (which has higher SNR and benefits short ATT brain territories) with late sampling (which has lower SNR and benefits long ATT territories). So far, this optimisation framework has only been applied to sequential multi-PLD PCASL protocols with a fixed and unoptimised label duration.

Time-encoding of the PCASL preparation using a Hadamard encoding scheme has been proposed as a more efficient method for acquiring multi-PLD/LD ASL data, due to the noise averaging that occurs during the decoding process (Dai et al., 2013; Günther, 2007; Wells et al., 2010). However, this reduced noise may be counteracted by reduced ASL signal due to the use of shorter LDs for each sub-bolus (Guo et al., 2018). Multiple variations of the time-encoded technique have been proposed in order to improve the SNR across the different time points (Teeuwisse et al., 2014), but so far the CBF accuracy of only fixed-LD time-encoded protocols have been compared with single-PLD and sequential multi-PLD/LD protocols and these protocols were not first optimised for CBF accuracy (Dai et al., 2013; Guo et al., 2018; Johnston et al., 2015). Therefore, the results of these comparisons may simply reflect the chosen protocol timings rather than the ultimate accuracy of each technique.

In this work, we aimed to establish which PCASL approach can achieve the most accurate CBF measurements. We did this by investigating the relative CBF accuracy of a single-PLD protocol, a wide range of multi-timepoint PCASL protocol designs from the literature, and several novel protocol hybrid designs which are introduced in this study (Figure 1). We first applied a previously developed optimisation framework (Woods et al., 2019) to the multi-timepoint protocol timings to ensure each protocol would optimally estimate CBF across an expected range of ATTs for healthy grey matter (GM) given the design constraints of each protocol. The CBF accuracy of these optimised protocols were then compared using Monte Carlo (MC) simulations, with a subset of protocols being compared in vivo.

**Figure 1:**
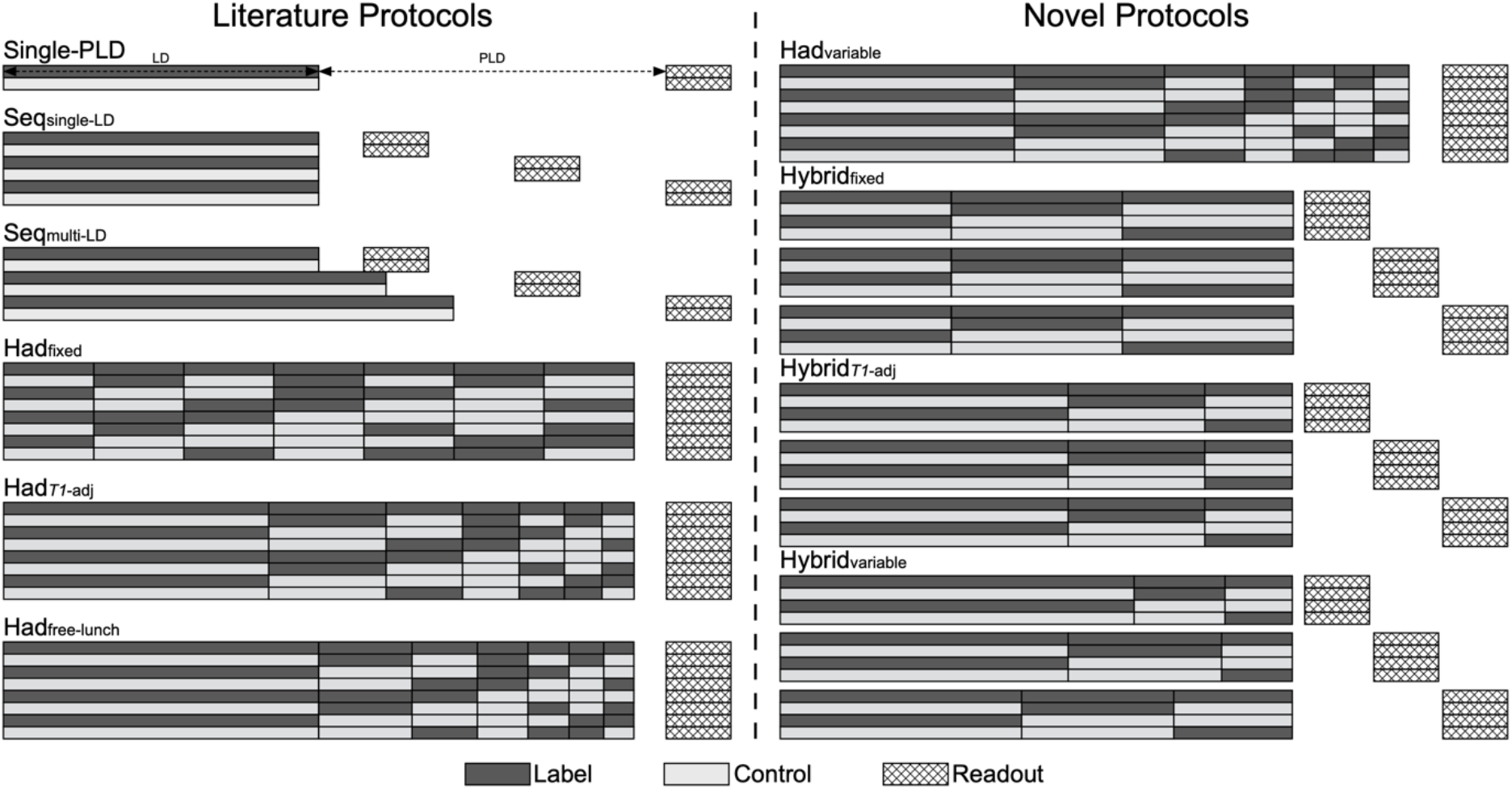
Example timing schematics of the PCASL label/control protocols used in this work. The number of label/control pairs and the size of the time-encoding matrices were optimised in each case; see text for details.

## 3. Theory

### 3.1. Literature protocol designs

The range of protocol designs investigated in this work are shown in Figure 1. The single-PLD and sequential multi-PLD, with a fixed LD, (Seq_single-LD_) protocol designs have been widely used in the literature to estimate CBF only or both CBF and ATT, respectively (Alsop et al., 2015; Alsop and Detre, 1996; Buxton et al., 1998; Dai et al., 2017; Gonzalez-At et al., 2000; Okell et al., 2013). Borogovac et al. (2010) suggested the use of multiple sequential LDs with a fixed PLD as a more SNR-efficient method for estimating CBF and ATT than fixed-LD multi-PLD methods, though this hypothesis was not tested. (Johnston et al., 2015) later demonstrated the use of both varying LDs and PLDs (referred to here as Seq_multi-LD_) to estimate CBF and ATT, but this implementation did not use inversion pulses for background suppression (BGS), instead relying only on pre-saturation to facilitate *T_1_* estimation from the ASL data, which may have affected the resulting CBF accuracy. In this study, we investigated both Seq_single-LD_ (a single fixed LD with *N* PLDs) and Seq_multi-LD_ protocols (*N* LDs with *N* PLDs).

Günther (2007) introduced time-encoded PCASL as an efficient method for generating multi-timepoint ASL data. The PCASL pulse train is split into *M* sub-boluses which vary between label and control conditions within each TR according to a predesigned encoding matrix (a Hadamard matrix being the most efficient encoding). The acquired data is then decoded using the same encoding matrix generating *M* perfusion weighted images which reflect the effective LD and PLD of each sub-bolus; for a Hadamard encoding of size (*M* + 1) × *M* this results in a 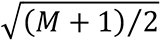 decrease in noise SD (assuming additive white Gaussian noise) and a scan time reduction of (*M* + 1)⁄(2 · *M*) compared to a matched timing sequential control - tag experiment (Dai et al., 2013).

The original time-encoded protocol used a fixed LD for all sub-boluses (Had_fixed_). Several variations were introduced by (Teeuwisse et al., 2014), including the free-lunch (Had_free-lunch_) and *T_1_*-adjusted (Had*_T1_*_-adj_) protocols. In the *T_1_*-adjusted protocol, the encoded LDs are set such that the total ASL signal originating from each sub-bolus is equal at the time of acquisition, thus accounting for the increased *T_1_* decay experienced by earlier sub-boluses and so maintaining an approximately constant level of SNR after complete bolus arrival. The free-lunch protocol uses the same long LD and PLD for the first encoded bolus as a typical single-PLD protocol, with the remaining sub-boluses filling this long PLD. After decoding, similar data to the single-PLD experiment is generated from the first sub-bolus, with the remaining sub-boluses generating extra temporal data without an increase in scan time. Figure 1 shows Had_free-lunch_ with *T_1_*-adjusted LDs, but any scheme may be used.

### 3.2. Hybrid protocol designs

Here, we introduce a novel protocol design which is a hybrid of the time-encoded and sequential protocols. Rather than using a fixed final PLD after the time-encoded preparation and acquiring multiple averages, there are *N* final PLDs which sequentially vary for each repeat of the same encoding matrix, allowing increased flexibility of the decoded timepoints. This results in *N* · *M* decoded timepoints (*N* final PLDs, *M* time-encoded sub-boluses). This design can trade-off the superior noise averaging of the time-encoding methods (larger encoding matrices result in more signal averaging) and the increased signal accumulation from longer LDs (typically achievable with smaller encoding matrices). We investigated the use of both fixed (Hybrid_fixed_) and *T_1_*-adjusted (Hybrid*_T1_*_-adj_) time-encoded LDs with this protocol design. The Hybrid protocols were previously presented in abstract form (Woods et al., 2018).

### 3.3. Variable-LD time-encoded and hybrid designs

The time-encoded and hybrid protocols do not have to be restricted to the designs discussed above, i.e. fixed and *T_1_*-adjusted LDs. It is possible for the individual encoded LDs and final PLDs to be chosen arbitrarily. As an extension to the comparison of the protocols detailed above, we tested whether there is a more optimal time-encoded LD pattern than the existing literature designs by optimising a time-encoded protocol and a hybrid protocol where each LD in the encoding matrix could be adjusted separately, rather than according to a predefined pattern. To increase the flexibility of the hybrid protocol even further, each of the *N* final PLDs was associated with a separate encoding matrix of *M* LDs, rather than repeating the same encoding matrix timings for each of the PLDs, leading to *N* · *M* decoded timepoints with *N* · *M* separate LD and PLD pairs. These protocols are referred to as Had_variable_ and Hybrid_variable_.

## 4. Material and methods

All optimisations, simulations and analysis, except CBF and ATT estimation, were performed using MATLAB (The MathWorks, Natick, MA).

### 4.1. Protocol optimisation

The multi-timepoint protocols described above were optimised for CBF accuracy, while treating ATT as a potentially confounding parameter, using a recently developed framework (Woods et al., 2019). The original implementation of the optimisation algorithm iterated through each of the *N* PLDs of a multi-PLD protocol, and for each, performed a grid search for the PLD value which minimised the mean Cramér-Rao Lower Bound (CRLB) variance across ATTs, taking into account the number of averages realisable in a given scan time. The principal of the optimisation for each protocol considered in this work was the same, but due to the different sizes of the timing parameter spaces, the implementation was adapted in each case, as described in Supporting information text 1 - Protocol optimisation. For each protocol, the number of effective PLDs, *N_T_*, was optimised for by running the optimisation for a range of *N_T_* and selecting the protocol with the minimum cost. *N_T_* was constrained to ≤15 to ensure multiple averages at each PLD.

The optimisation used a uniform ATT prior probability distribution with a representative GM range of 0.5 - 2 s for healthy volunteers (Alsop et al., 2015; Dai et al., 2017; Guo et al., 2018; Woods et al., 2019), sampled at 1 ms increments, with a 0.3 s linearly decreasing weighting beyond either end of the range to reduce edge effects. Since the optimisation does not depend on CBF (Woods et al., 2019), a CBF point prior of 50 mL/100g/min was used. The LD update grid searches were restricted to 0.1 s ≤ LD ≤ 1.8 s with 25 ms increments, ensuring the minimum LD was greater than 100 ms, as suggested by (Teeuwisse et al., 2014), with the longest LD matching the recommended single-PLD LD of 1.8 s (Alsop et al., 2015). The PLD update grid was 0.075 s ≤ PLD ≤ 2.3 s with 25 ms increments. Other settings included: single-shot readout with 638 ms of non-ASL time per TR (presaturation and readout); variable minimum TR (Wang et al., 2013) (where the TR is minimised for each timepoint); 5 minute scan duration. The CRLB was calculated using the standard CASL kinetic model from (Buxton et al., 1998), using the parameters in Table 1, assumed additive white Gaussian noise, as described in (Woods et al., 2019). The noise magnitude was calculated from preliminary in vivo data (noise SD of label and control data = 1.3 × 10^−3^ relative to *M_0_*).

**Table 1:** Model and sequence parameters used in the optimisations, Monte Carlo simulations and in vivo experiments.

| Parameter | Value |
| --- | --- |
| **Model** |  |
| $T_1$ of blood ($T_{1b}$) | 1.65 s (Lu et al. 2004) |
| $T_1$ of tissue ($T_{1t}$) | 1.445 s (Lin et al. 2001) |
| Labeling efficiency ($\alpha$) | 0.85 (Dai et al. 2008) |
| Brain/blood water partition coefficient ($\lambda$) | 0.9 mL/g (Herscovitch et al. 1985) |
| **Sequence** |  |
| RF labeling pulse duration | 500 μs duration (Gaussian) |
| RF labeling pulse interval | 1 ms |
| RF labeling flip angle | 20° |
| Mean labeling gradient | 0.8 mT/m |
| Gradient during labeling pulses | 6 mT/m |
| **Analysis** |  |
| CBF prior | $0 \pm 10^6$ mL/100g/min |
| ATT prior | $1.3 \pm 10^6$ s |

### 4.2. Monte Carlo simulations

Monte Carlo simulations were performed to investigate the performance of the optimised protocols under ideal conditions where the ground truth is known. Simulated data were generated for each protocol using the standard CASL kinetic model (Buxton et al., 1998) with the parameters in Table 1 for ATTs between 0.5 - 2 s at 0.01 s increments. White Gaussian noise was added to 2000 replicas of the label and control (or encoded) data at each ATT sample, using the same noise magnitude as the protocol optimisations above. The noisy difference data at each timepoint was then decoded according to the encoding scheme for each protocol. The data were then fit, and the estimates compared, as described below.

### 4.3. In vivo experiments

#### 4.3.1. Acquisition

To investigate the relative performance of the protocols given in Table 2 in vivo, 10 healthy volunteers (5 female, mean age 30.7, range 24 - 40) were recruited and scanned under a technical development protocol, agreed with local ethics and institutional committees, on a 3T Prisma system (Siemens Healthcare, Erlangen, Germany) with a 32-channel receive-only head coil. The Had_variable_ and Hybrid_variable_ protocols were not compared in vivo since they only led to marginal improvements in CBF accuracy during simulation (see Results 5.9). All scanning occurred during a single session for each subject (total scan duration ∼50 minutes). Volunteers were asked to lie still and stay awake throughout the scan. A nature documentary was shown to help maintain alertness.

**Table 2:**
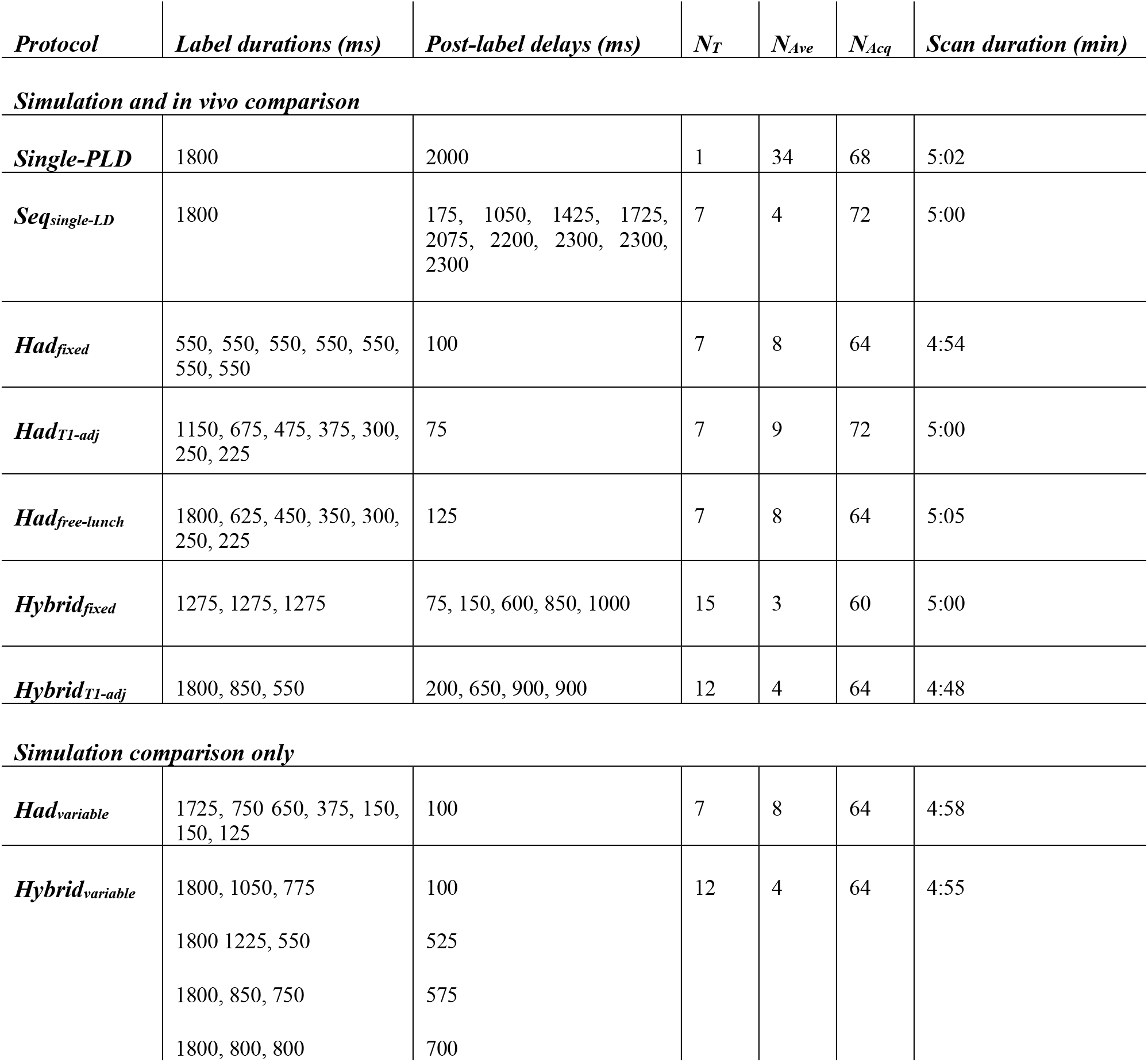
The optimised protocol timings for the protocols compared in vivo and in simulation. For the time-encoded (Had) and hybrid protocols, the LDs are given in chronological order and the number of LDs defines the size of the Hadamard encoding matrix used. For the Hybrid_variable_ protocol, each PLD is associated with the LDs on the same row. N_T_ is the number of effective PLDs, N_Ave_ is the number of averages, and N_acq_ is the number of acquired volumes for each scan.

The scan protocol included a 3-plane localiser and a 3D single-slab TOF angiography sequence used to position the PCASL labelling plane. A 3D *T_1_*-weighted MPRAGE sequence (1.5×1.5×1.5 mm^3^) was acquired for generating the brain and grey matter (GM) masks. Four calibration images were acquired with the same readout module as the PCASL data (see below) but with alternating in-plane phase encode direction to correct off-resonance distortions. Finally, the ASL scans were acquired in a pseudo-randomly permuted order for each subject to reduce the impact of physiological drift.

ASL imaging parameters were: single-shot 3D gradient and spin-echo (GRASE) readout (Feinberg and Oshio, 1991; Günther et al., 2005), TE 28.5 ms, variable minimum TR, excitation flip-angle 90°, refocussing flip-angle 120° (He et al., 2018; von Samson-Himmelstjerna et al., 2016), FOV 230×168×100 mm^3^, matrix 64×46×20, 20 acquired partitions, no parallel imaging acceleration, no slice-oversampling, centric partition ordering, bandwidth 2298 Hz/px, total readout duration 583 ms, spectrally-selective fat saturation. The imaging slab was placed in the transverse plane with the superior edge flush with the top of the brain. The excitation and refocussing pulse widths were 110 mm and 150 mm, respectively, to maximise the signal level within the nominal slab. Outer-volume suppression (OVS), using a cosine-modulated water suppression enhanced through *T_1_* effects (WET) module (Golay et al., 2005; Ogg et al., 1994), was used to improve the slab profile, similar to (Günther et al., 2005). Readout, phase-encode, and 3D encode directions were anterior-posterior, right-left, and feet-head, respectively.

PCASL labelling was achieved using the parameters in Table 1 with the labelling plane positioned in the transverse plane bisecting the V3 section of the vertebral arteries (Okell et al., 2013). BGS was performed with a slab-selective WET presaturation module (Golay et al., 2005; Ogg et al., 1994) immediately before the start of labelling and two optimally timed slab-selective C-shaped FOCI pulses (µ = 1.5, β = 1349 s^−1^, *A_max_* = 20, duration 10.24 ms) (Ordidge et al., 1996; Payne and Leach, 1997). The presaturation and inversion slabs were parallel to the labelling plane and covered the entire brain, with the inferior edge at the centre of the labelling plane. For each protocol, the inversion pulses were timed to null two *T_1_* values (700 ms and 1400 ms) 100 ms before excitation using the formula in (Günther et al., 2005). The inversion pulses were interleaved with the PCASL labelling when the optimal inversion times occurred during the labelling period, as in (Dai et al., 2012, 2010), leading to more uniform BGS across a range of timings (Supporting Information Figure S1).

The calibration images were acquired using presaturation followed by a 10 s delay to allow controlled and near-complete magnetisation recovery before the 3D-GRASE readout.

#### 4.3.2. Preprocessing

Preprocessing of the in vivo data was performed using tools from the FSL toolbox (Jenkinson et al., 2012). The ASL data were motion-corrected and registered to the mean calibration data with rigid-body registration using FLIRT (Jenkinson, 2002; Jenkinson and Smith, 2001), before correcting for susceptibility induced off-resonance geometric distortions using TOPUP (Andersson et al., 2003). Brain and GM masks were generated from the structural data using BET (Smith, 2002) and FAST (Zhang et al., 2001). These were transformed to ASL space after image registration (Greve and Fischl, 2009) and had thresholds applied (brain mask 90%, GM mask 50% tissue partial volume).

The edges of the brain-masked calibration image were eroded before being expanded using a mean filter and brain masked again to remove low-intensity voxels at the edge of the brain which can lead to erroneously high CBF values during the voxelwise calibration step. It was then smoothed (Gaussian kernel, σ = 2.5 mm) to improve SNR, as recommended (Alsop et al., 2015).

The perfusion-weighted images were generated by pairwise subtracting or decoding the preprocessed ASL images. They were then calibrated prior to fitting to account for scaling factors by voxelwise dividing by the smoothed calibration image and the labelling efficiency and multiplying by the blood–brain partition coefficient (Table 1).

### 4.4. Model fitting

CBF and ATT were estimated identically for the simulated data and in vivo data using the variational Bayesian inference algorithm, BASIL (Chappell et al., 2009). In each voxel, this approach not only provides estimates of CBF and ATT but also uncertainty estimates in the form of the standard deviation of the marginal posterior distributions. The standard CASL kinetic model (Buxton et al., 1998) was used with the parameters in Table 1. Fitting was initialised with a coarse grid search for robustness (bounded by 0 ≤ CBF ≤ 200 mL/100g/min and 0 ≤ ATT ≤ 2.5 s, sampled every 1 mL/100g/min and 0.01 s). The BASIL fitting priors were noninformative to minimise bias in the resulting parameter estimates. Negative CBF and ATT estimates were set equal to zero. The single-PLD data was only fit for CBF with the ATT fixed at 1.3 s; this value was found to minimise the theoretical CBF bias across the ATT range 0.5 - 2 s. The data was not averaged before fitting.

In vivo ground truth CBF and ATT estimates were generated by fitting the combined data from all protocols, similar to (Woods et al., 2019). To account for the different noise levels between protocols, BASIL was given 3 noise magnitudes to estimate in an approach similar to weighted NLLS fitting (Chappell et al., 2009). Three noise magnitudes were used because there were 3 categories of data with similar noise magnitudes after decoding: the non-time-encoded data (single-PLD and sequential protocols), the 8×7 Hadamard encoded protocols, and the 4×3 Hadamard encoded hybrid protocols (see Results 5.1). To investigate whether these ground truth estimates were biased towards certain protocols and whether modelling the 3 noise magnitudes is beneficial, ground truth estimates for the MC simulation data were identically generated with either 1 or 3 noise magnitudes.

### 4.5. Comparison metrics

The CBF estimates of each protocol were compared in three different ways for both simulation and in vivo data: (1) the marginal posterior probability distribution SDs output by BASIL were used as a measure of uncertainty in the CBF estimates (Chappell et al., 2009), and are sensitive to how well the kinetic model fits the data; (2) the root-mean-squared-error (RMSE) relative to the ground truth estimates were used as a measure of accuracy, incorporating both systematic bias and noise contributions, similar to (Woods et al., 2019); and (3) the test-retest RMSE for each scan was calculated by splitting the data into two 2.5 minute data sets and separately fitting each half, giving a measure of within-session repeatability, which is independent of any ground truth estimates or uncertainties derived from the fitting process. Note, for (3) the Had*_T1_*_-adj_ data were split into the first 4 and last 5 averages while the Hybrid_fixed_ errors could not be calculated because there were only 3 averages (see Table 2).

### 4.6. In vivo analysis

Only voxels within the GM masks were used in the analysis. To eliminate poorly fit ground truth data from the analysis, voxels with ground truth posterior SDs more than 3 times the inter-quartile range above the 75th percentile for either CBF or ATT were excluded (Tukey, 1977). This resulted in upper bounds on the ground truth posterior SDs of 2.9 mL/100g/min and 0.061 s. Voxels were also excluded if the posterior SDs for any individual protocol were > 500 mL/100g/min or > 50 s, which would suggest an extremely poor fit, perhaps arising from motion or other artefacts, and could bias the resulting comparison. This extremely poor fit criteria was also used for the MC simulation analysis.

The comparison metrics were calculated on a voxelwise and subjectwise basis. Standard errors for the voxelwise metrics were calculated by bootstrap sampling (Efron, 1979) across the 10 subjects using 1000 samples, where the relevant statistical measure (mean SD, RMSE, or test-retest RMSE) was performed on each bootstrap sample. Each sample is a selection of 10 randomly chosen subjects, selected with replacement, meaning a given sample could contain multiple copies of the same subject’s data. The SDs generated from these bootstrap distributions reflect the variability in the voxelwise metrics due to the sampled subjects. This approach gives a more conservative standard error than would be calculated from the combined voxelwise data across subjects due to the large number of voxels.

## 5. Results

### 5.1. Optimised protocols

The optimised timings for each protocol are shown in Table 2 and the predicted CBF SDs (CRLBs) are shown as a function of ATT in Figure 3. The results of the Had_variable_ and Hybrid_variable_ protocols are reported separately in Results 5.9 due to the marginal improvements achieved with these protocols.

**Figure 3:**
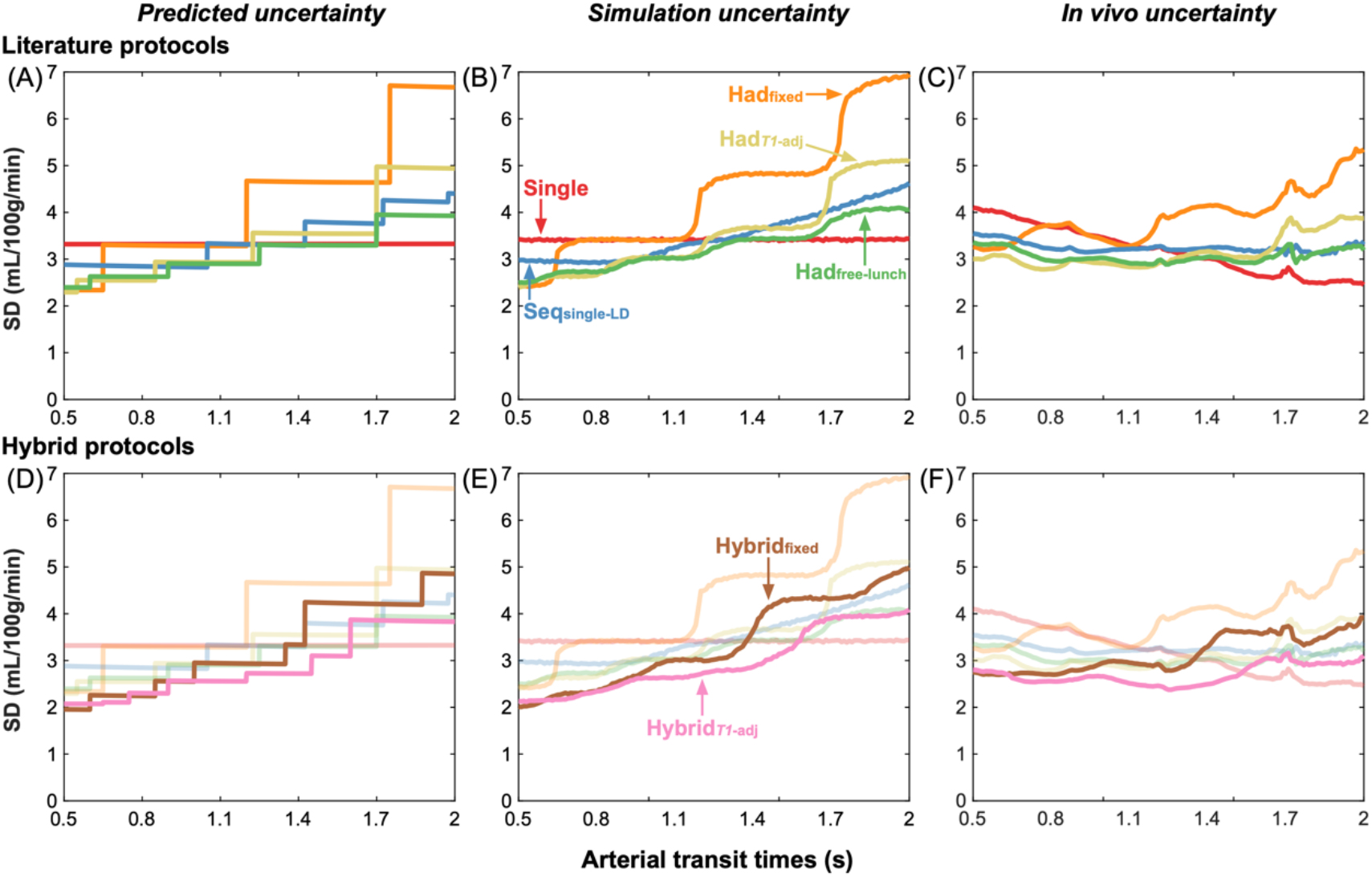
The predicted (Cramer-Rao lower bound SDs) (A,D), simulation (Monte Carlo simulation posterior SDs) (B,E), and in vivo (posterior SDs) (C,F) CBF uncertainty measures for the literature protocols (top) and the proposed hybrid protocols (bottom) shown across ATTs. For the simulation and in vivo results, the median SD at each ATT is plotted. A sliding window was used to plot the in vivo data with window size 0.1 s and step size 0.01 s.

The increasing density of PLDs at later times in both sequential protocols (Seq_single-LD_ and Seq_multi-LD_ - see Supplementary Table X) is similar to the CBF optimised multi-PLD protocol in (Woods et al., 2019) but differs here because a different ATT range and a 3D readout were used. Due to the similarity with the Seq_single-LD_ protocol timings, and the marginal improvement in predicted CBF errors, Seq_multi-LD_ was not used in further comparisons. A 4×3 encoding came out as optimal for the Had_fixed_ protocol (15% lower average CBF CRLB SD than 8×7, see Supporting Information Figure S8 and Supporting Information Table S1), but due to the more common use of the 8×7 encoding and the large jump in the CBF error part way through the ATT range, the 8×7 protocol was used in further comparisons. The optimal Had_free-lunch_ protocol was an 8×7 encoding with 6 *T_1_*-adjusted LDs.

### 5.2. In vivo CBF and ATT maps

Figure 2 shows the spatial maps of the CBF and ATT estimates, their uncertainties (expressed as the SD of the posterior distribution), and the errors relative to the ground truth estimates for each tested protocol for a single representative subject. The CBF and ATT maps are shown for all subjects in Supporting Information Figure S2 and Supporting Information Figure S3. There is good agreement in broad spatial variations of both CBF and ATT between the protocols, demonstrating the overall consistency of the estimates. However, the error maps highlight the variation between protocols in over/under-estimating CBF and ATT. Particularly evident, is the effect that the assumed single-PLD ATT had on the single-PLD CBF errors: regions where the assumed ATT was an overestimate led to the CBF being underestimated, relative to the ground truth estimates.

**Figure 2:**
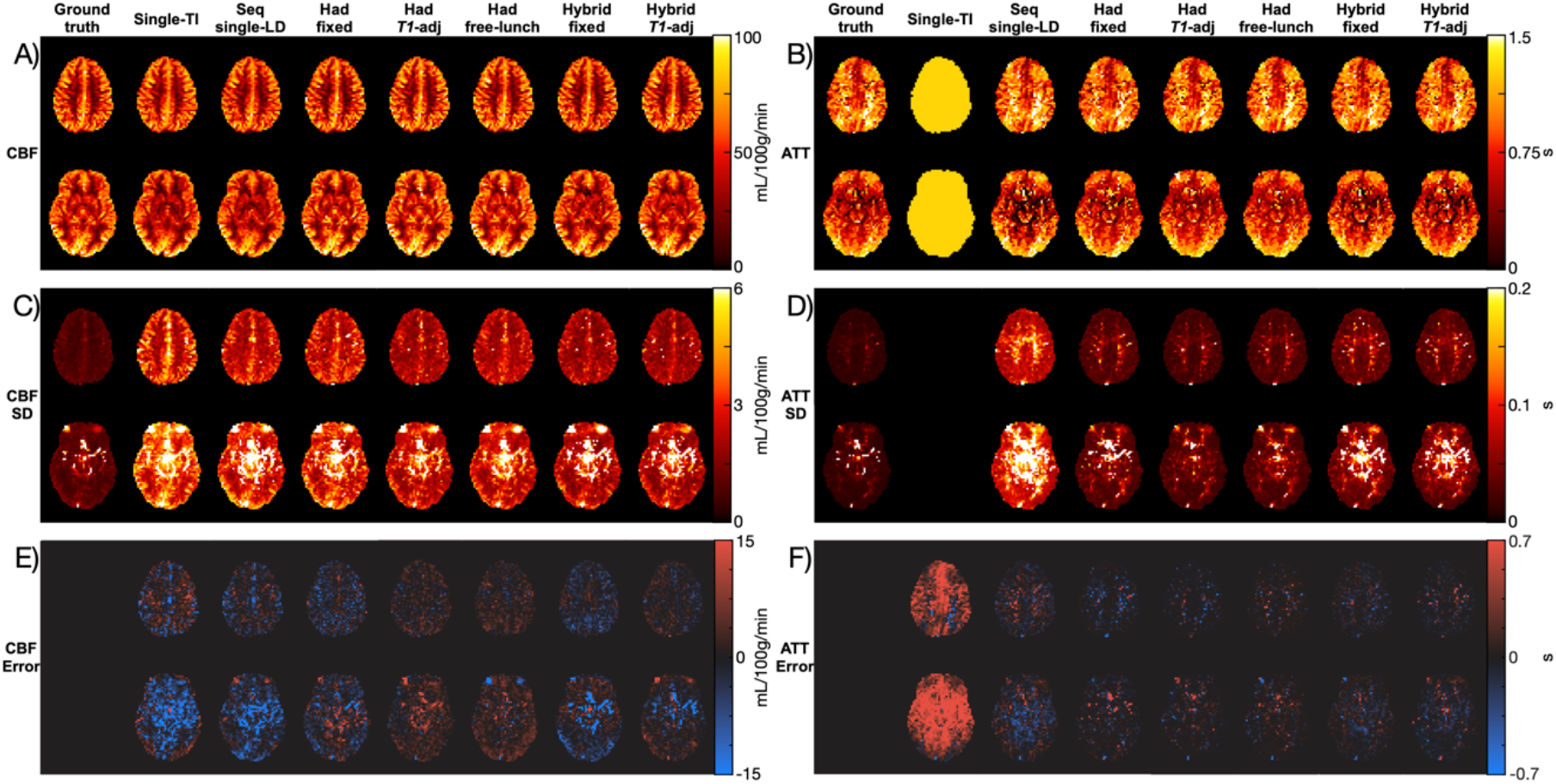
The CBF (A, C, E) and ATT (B, D, F) mean estimates (A, B), uncertainties (C, D), expressed as the marginal posterior probability distribution output by BASIL, and the errors relative to the ground truth mean estimates (E, F) shown as protocol estimates - ground truth estimates. Two slices from a representative subject are shown. The colour maps use perceptually uniform colour maps, developed by (Kovesi, 2015).

High uncertainties in the lower slice of the CBF and ATT SD maps can be seen in regions consistent with the known location of large arteries. Due to the presence of these elevated SDs in the single-PLD data, which has a long PLD of 2 s, it was assumed to be largely caused by signal dephasing of pulsatile flow during the GRASE readout, rather than macrovascular ASL signal. However, these large arteries may contain residual ASL signal for the protocols with short PLDs. High uncertainties can also be seen in the sagittal sinus and from eye motion. These voxels were not included in the quantitative comparisons (see Results 5.3).

### 5.3. In vivo data selection

There were a total of 79,211 voxels in the GM masks across all 10 subjects. Of these, 6.2% were excluded due to poor ground truth CBF and ATT fits (posterior SDs > 2.9 mL/100g/min or > 0.061 s) and a further 4.1% were excluded because there were extremely poor fits in one or more of the individual scans (posterior SDs > 500 mL/100g/min or > 50 s).

Of the included voxels, 90% of the ground truth ATTs lay between 0.5 - 1.51 s (5th - 95th percentiles, median = 0.97 s). Supporting Information Figure S4 shows, for a single subject, that the excluded voxels are mostly located where one would expect large arteries to be. For the included voxels, the mean grey matter CBF estimates were not significantly different across protocols on the subject level (Wilcoxon signed rank test), averaging at 57.17 ± 0.48 mL/100g/min (mean ± SD across protocols).

### 5.4. Trends across ATTs

The predicted CBF uncertainties (the CRLB SDs) for the literature protocols and novel hybrid protocols are shown in Figure 3(A, D) as a function of ATT for a fixed CBF of 50 mL/100g/min. The single-PLD CBF uncertainties were flat across the ATT range because it is only dependent on the noise magnitude, which is assumed to be constant across all ATTs. The sharp changes in uncertainties across ATTs for the multi-timepoint protocols are where ATT=PLD or ATT=LD+PLD for one or more of the LD/PLD pairs. As the ATT increases, these discontinuities represent the transition of a data point to either no longer sampling the inflow section of the kinetic model (LD+PLD<ATT) or moving from the tracer decay portion of the model (ATT<PLD) to the inflow portion (ATT<LD+PLD<LD+ATT). Both cases result in an increase in the CBF uncertainty.

Of the literature protocols, Had_free-lunch_ maintained the lowest uncertainties across most of the ATT range. Had*_T1_*_-adj_ performed similarly to Had_free-lunch_ at short ATTs, reflecting the similarity in the timings of their last 6 time-encoded LDs, but had much larger uncertainties at ATT > 1.7. Seq_single-LD_ maintained similar uncertainties across the ATT range to Had_free-lunch_ and Had*_T1_*_-adj_. Had_fixed_ had the highest predicted CBF uncertainties across most of the ATT range. All the multi-timepoint protocols had reduced uncertainties at short ATTs compared to the single-PLD protocol but were worse at longer ATTs.

Both hybrid protocols achieved lower predicted CBF uncertainties at almost all ATTs relative to their non-hybrid analogues. The Hybrid*_T1_*_-adj_ protocol also maintained a lower uncertainties than the other multi-timepoint protocols at almost all ATTs and had lower uncertainties than the single-PLD protocol for most of the distribution.

The median CBF uncertainties from the MC simulations (the marginal posterior probability distribution SDs from the Bayesian fitting) are shown in Figure 3(B, E) and follow the trends of the predicted uncertainties extremely closely, validating the expected performance of each protocol under ideal conditions. The CBF uncertainty discontinuities are visible but are more gradual due to the blurring effect of noise on ATT estimation.

The in vivo median CBF posterior uncertainties (Figure 3(C, F)) exhibit similar relative performance for each protocol, but there is a general decrease in the uncertainties at longer ATTs for all protocols compared to the predicted and simulation CBF uncertainties. This is thought to be due to the correlation between ATT and receive coil SNR (shorter ATTs are generally found closer to the centre of the brain where the SNR is lower - see Discussion 6.3). Similar jumps in the uncertainties can be seen, especially for the Had_fixed_ protocol.

### 5.5. Uncertainty: mean posterior distribution SD

The mean MC simulation and in vivo voxelwise CBF posterior SDs across all ATTs, which represents the average uncertainty in the CBF estimates, are shown in Figure 4. The simulation results are shown with both the uniform ATT distribution and weighted by the measured in vivo ATT distribution.

**Figure 4:**
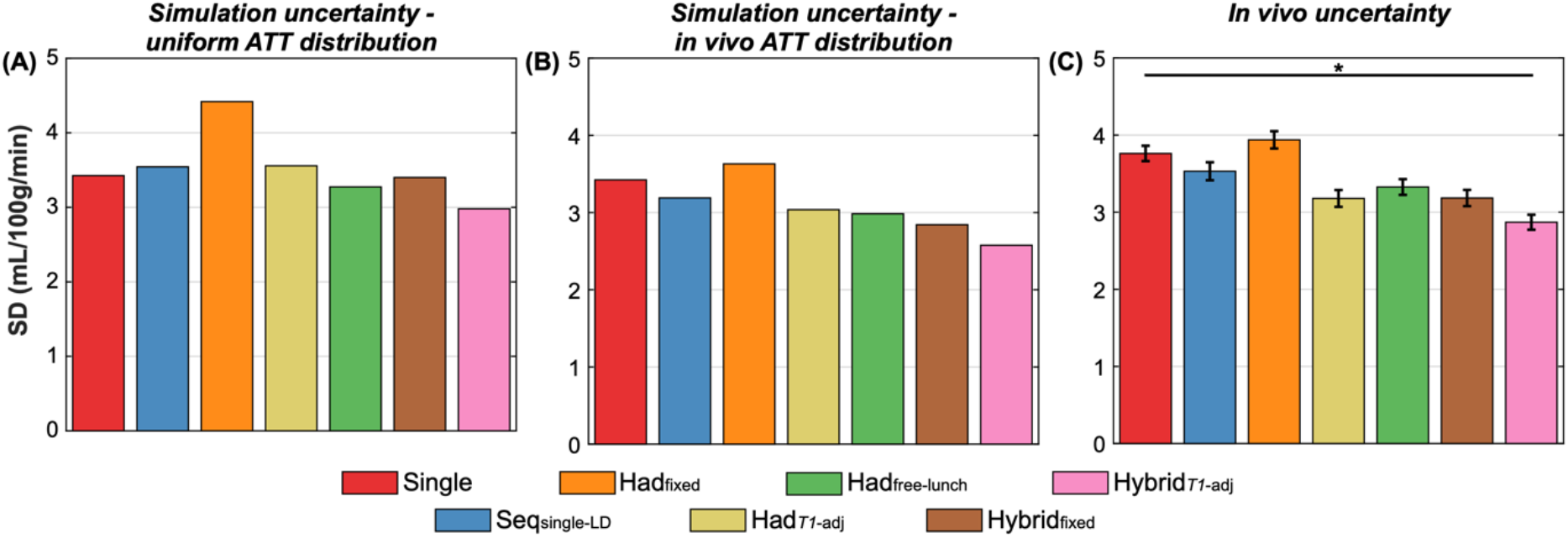
The simulation (A, B) and in vivo (C) mean posterior SDs across all voxels. (A) shows the simulation results for the uniform ATT distribution, while (B) shows the simulation results weighted by the measured in vivo ground truth ATT distribution. In vivo, the means and standard errors of the bootstrap distributions are shown (see methods). All differences were significant (two-sided paired Wilcoxon signed-rank test, Bonferroni correction for 6 comparisons, α<0.05).

Of the literature protocols, Had_free-lunch_ had the lowest simulation CBF uncertainty across both the uniform and the in vivo ATT distributions, including single-PLD (4% lower for the uniform ATT distribution). Across all the protocols, Hybrid*_T1_*_-adj_ had the lowest simulation CBF uncertainty (13% and 9% lower mean posterior SD than single-PLD and Had_free-lunch_, respectively, for the uniform ATT distribution).

The in vivo results are similar to the uniform ATT distribution simulation results but much more closely match the simulation uncertainties when they are weighted by the in vivo ATT distribution. The upweighting of shorter ATTs found in vivo led to several differences including single-PLD having worse CBF uncertainty than all protocols except Had_fixed_ and the performance of Had*_T1_*_-adj_ being improved relative to Had_free-lunch_ and Seq_single-LD_. In vivo, Had*_T1_*_-adj_ had the lowest average CBF uncertainty of the literature protocols (16% and 4% lower mean posterior SD than single-PLD and Had_free-lunch_, respectively), while Hybrid*_T1_*_-adj_ maintained the lowest average CBF uncertainty of all the protocols in all cases (24% and 14% lower than single-PLD and Had_free-lunch_, respectively, in vivo).

The subjectwise data for all three comparison metrics are shown in Supporting Information Figure S5 and demonstrate similar trends to the voxelwise comparisons, though with fewer significant differences between protocols due to the lower statistical power of these comparisons.

### 5.6. Accuracy: RMSE relative to ground truth

Figure 5 shows the simulation and in vivo voxelwise RMSEs, which represents a measure of accuracy in the CBF estimates, including both systematic bias and precision, with a lower RMSE meaning a protocol was more accurate. As for the posterior SDs, Had_free-lunch_ had the best simulation CBF accuracy of the literature protocols in the simulations (18% lower RMSE than single-PLD), but Had*_T1_*_-adj_ had the best accuracy in vivo (40% and 5% lower RMSE than single-PLD and Had_free-lunch_, respectively). Over all the protocols, Hybrid*_T1_*_-adj_ had the best CBF accuracy in both simulation (24% and 7% lower RMSE than single-PLD and Had_free-lunch_, respectively, for the uniform ATT distribution) and in vivo (47% and 15% lower RMSE than single-PLD and Had_free-lunch_, respectively). The accuracy of single-PLD is poorer relative to the multi-timepoint protocols than in the uncertainty comparison due to the bias caused by assuming a fixed ATT, which is estimated in the ground truth data and multi-timepoint protocols.

**Figure 5:**
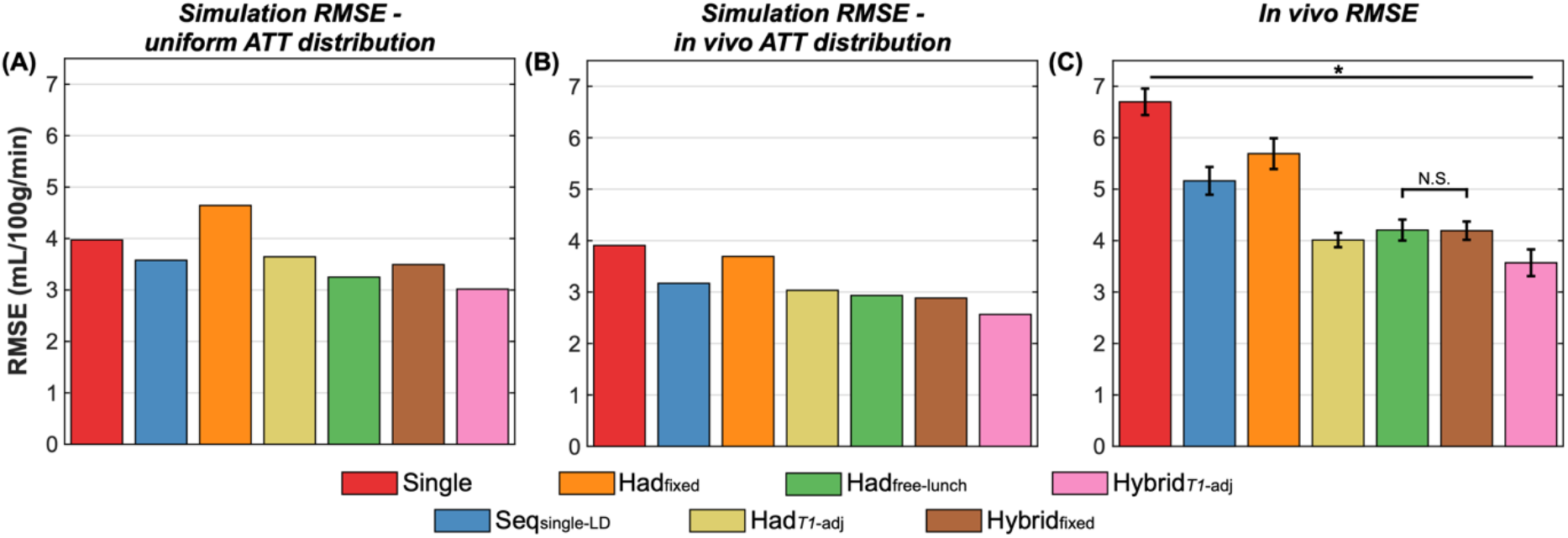
The simulation (A, B) and in vivo (C) RMSEs across all voxels. (A) shows the simulation results for the uniform ATT distribution, while (B) shows the simulation results weighted by the measured in vivo ground truth ATT distribution. In vivo, the means and standard errors of the bootstrap distributions are shown (see methods). All differences were significant except for Had_free-lunch_ vs Hybrid_fixed_ in vivo (two-sided paired Wilcoxon signed-rank test, Bonferroni correction for 6 comparisons, α<0.05).

Supporting Information Figure S6 demonstrates that the ground truth CBF estimates generated with one noise magnitude have a clear bias towards the single-PLD and Seq_single-LD_ protocols, whereas the ground truth estimates generated using three noise magnitudes underestimate the RMSEs for all protocols by a similar amount, effectively removing the relative bias between protocols in the comparison.

### 5.7. Repeatability: test-retest RMSE

The mean MC simulation and in vivo test-retest voxelwise CBF RMSEs are shown in Figure 6. A lower test-retest RMSE means a protocol was more repeatable. Note, a further 2.4% of the in vivo GM voxels were excluded from this comparison because one or more of the 2.5 minute scans had CBF or ATT posterior SDs > 500 mL/100g/min or > 50 s, suggesting very poor fits.

**Figure 6:**
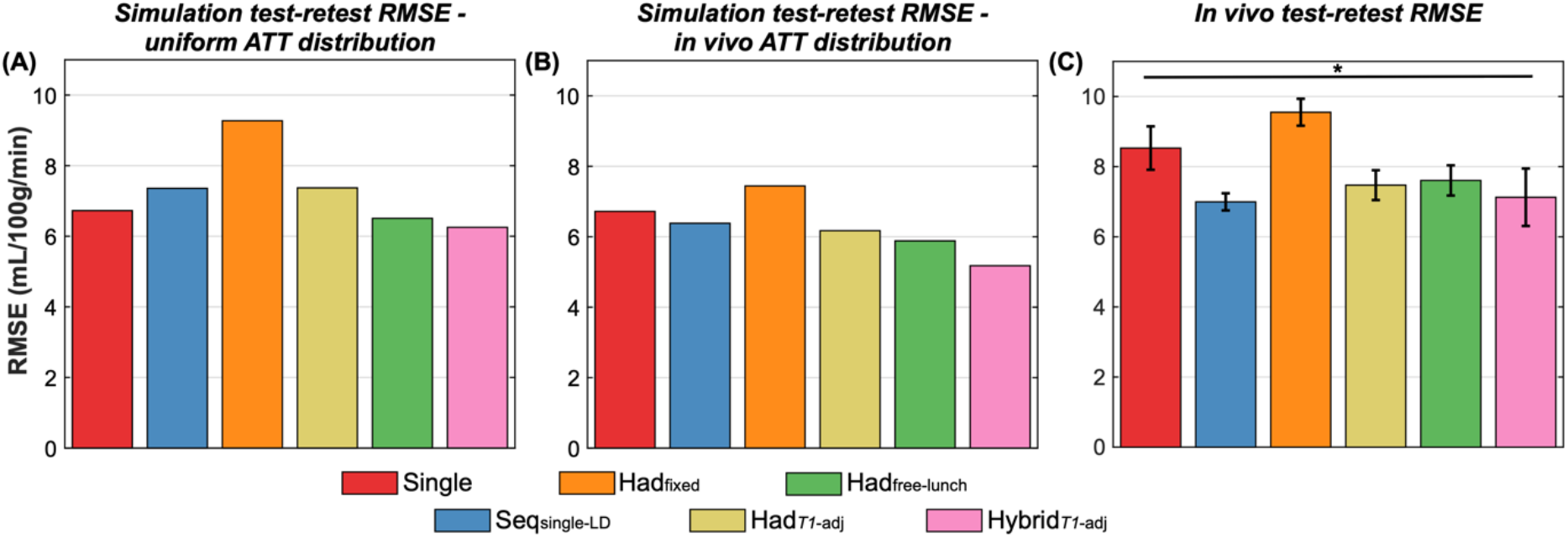
The simulation (A, B) and in vivo (C) test-retest RMSEs across all voxels. (A) shows the simulation results for the uniform ATT distribution, while (B) shows the simulation results weighted by the measured in vivo ground truth ATT distribution. In vivo, the means and standard errors of the bootstrap distributions are shown (see methods). All differences were significant (two-sided paired Wilcoxon signed-rank test, Bonferroni correction for 6 comparisons, α<0.05).

The simulation results reflect those of the uncertainty metric, with Had_free-lunch_ having the best repeatability of the literature protocols (3% lower test-retest RMSE than single-PLD for the uniform ATT distribution) while Hybrid*_T1_*_-adj_ had the best repeatability overall (7% and 4% lower test-retest RMSE than single-PLD and Had_free-lunch_, respectively, for the uniform ATT distribution). This again demonstrates that more robust CBF estimates can be obtained with certain multi-timepoint protocols than a single-PLD protocol, in this case using a metric which is not reliant on uncertainty estimates from the fitting algorithm nor any estimated ground truth.

As before, there were differences due to the shorter average ATTs seen in vivo than were simulated, causing the in vivo single-PLD CBF repeatability to be worse relative the multi-timepoint protocols. Another result also only seen in vivo was that Seq_single-LD_ had the best repeatability of all the protocols (RMSE = 7.00±0.24 mL/100g/min), better than Hybrid*_T1-adj_* (RMSE = 7.13±0.82 mL/100g/min). However, the subjectwise analysis, shown in Supporting Information Figure S5(C), demonstrates that there was one subject with much higher CBF test-retest RMSE for Hybrid*_T1_*_-adj_ than the other subjects. There was an average GM CBF increase of 10 mL/100g/min between the two halves of the Hybrid*_T1_*_-adj_ scan for this subject, possibly due to a change in subject alertness (Clement et al., 2018). After removing this subject from the comparison, Hybrid*_T1_*_-adj_ had the best CBF repeatability across all protocols (test-retest RMSE = 6.33±0.41 ml/100g/min: 28% and 15% lower than single-PLD and Had_free-lunch_, respectively) while the Seq_single-LD_ RMSE was relatively unaffected (7.01±0.27 mL/100g/min) (see Supporting Information Figure S9).

### 5.8. Arterial transit time

Although the protocols were not optimised for ATT accuracy, the results of the ATT comparisons are briefly described here. The in vivo voxelwise measures of ATT uncertainty, accuracy, and repeatability are shown in Supporting Information Figure S7 and demonstrate that the time-encoded and hybrid protocols all have more confident, accurate, and repeatable ATT estimates than Seq_single-LD_. Had*_T1_*_-adj_ had the lowest uncertainty, while Had*_T1_*_-adj_ and Hybrid*_T1_*_-adj_ both had the highest accuracy and best repeatability.

### 5.9. Had_variable_ and Hybrid_variable_ Protocols

The optimal Had_variable_ and Hybrid_variable_ protocol timings are given in Table 2 and the MC simulation uncertainties are shown in Figure 7. The Had_variable_ timings and uncertainties are similar to those of the Had_free-lunch_ protocol, though the average uncertainty is slightly lower for Had_variable_. Similarly, the optimised Hybrid_variable_ protocol only provided a small reduction in uncertainty relative to Hybrid*_T1_*_-adj_. These results suggest that the constraints of the Had_free-lunch_ (with *T_1_*-adjusted LDs) and Hybrid*_T1_*_-adj_ protocols are near optimal within their respective class of protocols, making them attractive protocol designs due to the reduced optimisation complexity resulting from their timing constraints. For these reasons, Had_variable_ and Hybrid_variable_ were not included during the in vivo comparison.

**Figure 7:**
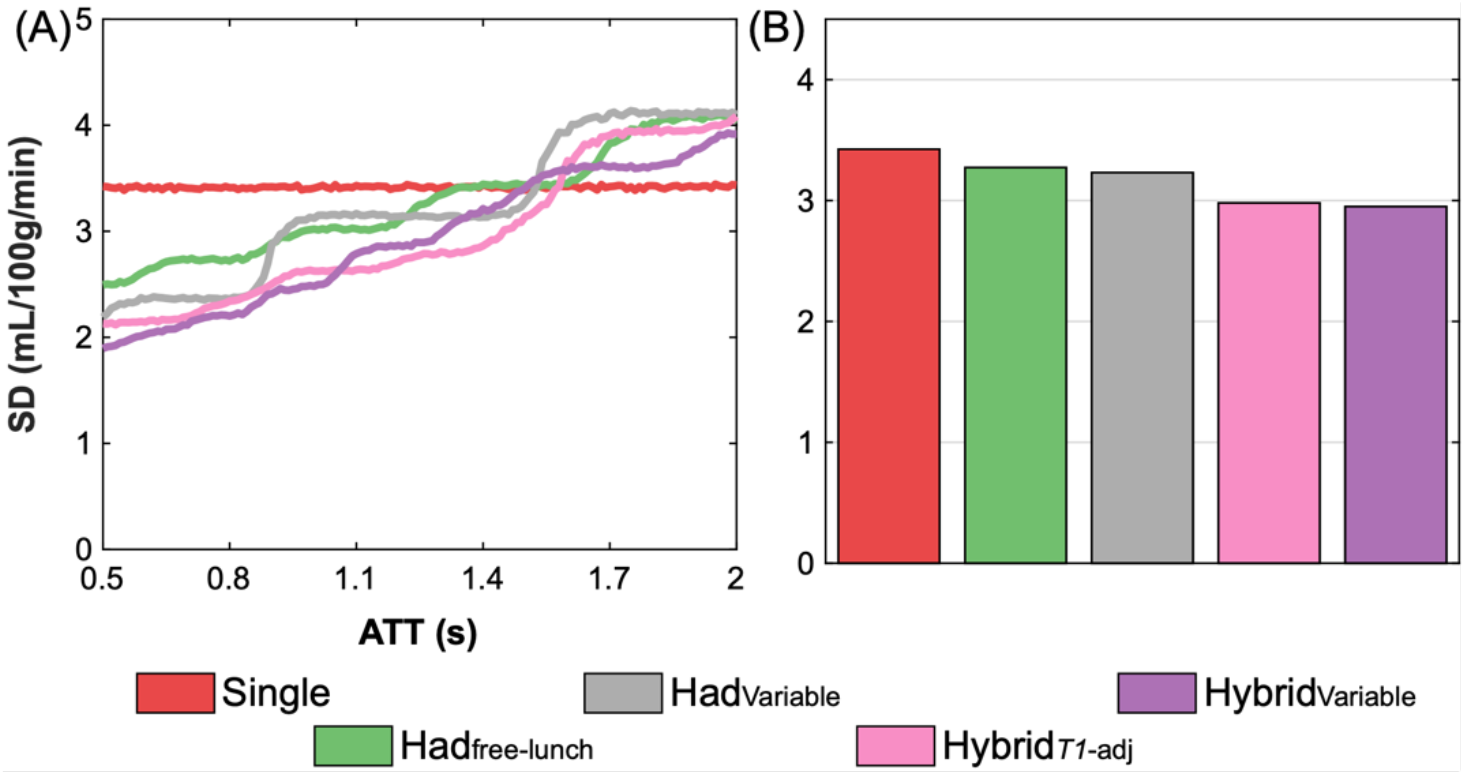
The MC simulation CBF posterior SDs (uncertainty) for Had_variable_, Hybrid_variable_, and a selection of previously compared protocols. (A) the median uncertainty for each protocol across ATTs, (B) the mean uncertainty for each protocol across the whole ATT range.

## 6. Discussion

In this study, a wide range of PCASL protocol designs were optimised for CBF accuracy, using a previously developed Cramér-Rao Lower Bound algorithm, and their CBF estimates compared using Monte Carlo simulations and in vivo experiments, which were in good agreement. The CBF estimates were compared with: (1) the standard deviation of the marginal posterior probability distributions from the fitting algorithm as a measure of uncertainty; (2) the RMSEs of the estimates relative to the ground truth estimates as a measure of accuracy, which includes both random variability and systematic biases; and (3) the RMSEs of the test-retest estimates as a measure of repeatability.

It was demonstrated that the Hybrid*_T1_*_-adj_ protocol had the most confident, most accurate and most repeatable CBF estimates of all the tested protocols, including the single-PLD protocol. This highlights the benefit of generating multi-timepoint ASL data from both time-encoded LDs and sequential PLDs. This hybrid method benefitted from the longer LDs possible with a smaller encoding matrix, but still achieved a time-decoding noise reduction factor of 2 and maintained a sufficiently well sampled range of unique PLDs due to the use of multiple sequential PLDs.

These results also highlight that, even though the multi-timepoint protocols have lower SNR at each timepoint compared to the single-PLD protocol, some can achieve more accurate CBF estimates on average across a range of ATTs. This is because the noise in multi-timepoint data is essentially averaged across the data during the fitting process, resulting in similar noise averaging to the single-PLD protocol, but with data that more effectively samples the signal curve across the range of ATTs.

Of the protocol designs from the literature, the Had_free-lunch_ design with *T_1_*-adjusted LDs was found to have CBF estimates that were more confident, accurate, and repeatable than the other literature designs for the uniform ATT distribution used in the simulations. Due to the shorter average ATTs witnessed in vivo, however, the Had*_T1_*_-adj_ protocol outperformed Had_free-lunch_. It was seen in simulation that Seq_single-LD_ produced similarly confident, accurate, and repeatable CBF estimates on average to Had*_T1_*_-adj_. This suggests that the averaging benefit from time-decoding for Had*_T1_*_-adj_ is similar to the benefit of longer LDs and more flexible PLDs for Seq_single-LD_. It is also apparent that the use of fixed-LDs in time-encoded PCASL is a sub-optimal design for CBF estimation.

### 6.1. Protocol optimisation

This study was restricted to protocols optimised solely for CBF accuracy. It is also possible to simultaneously optimise for CBF and ATT accuracy (Owen et al., 2016; Sanches et al., 2010; Santos et al., 2010; Woods et al., 2019; Xie et al., 2008) but we chose to focus on CBF estimation for two reasons: 1) CBF is often the main parameter of interest with knowledge of ATT predominantly being used to correct ATT related biases in the CBF estimates, and 2) optimising for only one parameter makes interpretation of the final protocols and their relative parameter estimation accuracy simpler. However, there is nothing to prevent the optimisation framework being used to also, or solely, optimise for ATT accuracy (Woods et al., 2019).

The optimised Seq_multi-PLD_ protocol included only one LD shorter than 1.8 s, suggesting that it is not optimal to use short LDs with short PLDs for CBF estimation for the investigated ATT range, a technique previously used in the literature (Johnston et al., 2015; Zhao et al., 2015). It also does not appear optimal to perform multi-timepoint acquisitions by only varying the LD (Borogovac et al., 2010).

The single-PLD protocol used in this study was not optimised using the CRLB framework, as used for the multi-timepoint protocols. Use of this framework would maximise the protocol’s average SNR, likely resulting in a PLD shorter than the longest expected ATT, which is in contrast with the recommended approach (Alsop et al., 2015) and could have resulted in potentially large CBF underestimation in regions where ATT>PLD (Guo et al., 2018). Future work could investigate the tradeoff between accuracy and precision of the single-PLD protocol with a shorter PLD in comparison to the best performing multi-timepoint protocols presented in this study.

The standard time-encoded protocols were relatively simple and fast to optimise, due to the reduced dimensionality of the timing parameter space enforced by the design constraints. This contrasts with the sequential and hybrid protocols which must be iteratively optimised, and therefore take more time; the Seq_mutli-LD_, Had_variable_, and Hybrid_variable_ protocols also required many random initialisations to avoid local minima. The standard time-encoded protocols might, therefore, make ideal candidates for real-time protocol optimisation since they can be quickly adjusted during a scan to better match patient specific ATT information generated from preceding TRs (Xie et al., 2010).

Hadamard-encoding schemes were used for the time-encoded protocols because these provide the most efficient encodings. However, they can only be of size (rows×columns) 2k×(2k-1), for k=1,2,4,6,8,10,…. Less efficient encodings may provide more flexibility in the protocol timings and could be explored with the same optimisation framework used in this work.

### 6.2. Choice of ATT prior

A uniform ATT prior distribution of 0.5 - 2 s was chosen based on the ATT range seen in (Woods et al., 2019), which used a similar labelling plane placement. However, the in vivo ATTs in this study were generally shorter, with 95% of the ground truth ATTs ≤1.51 s. This may be due in part to the use of a visual stimulus to maintain subject alertness, which can lead to a reduction in ATTs in the visual cortex (Qiu et al., 2010), a region which typically has longer ATTs than other GM brain regions (Dai et al., 2017). 5.2% of the voxels had ATTs <0.5 s, which was outside the optimised ATT range. However, if these voxels are excluded from the analysis, the results are similar and the conclusions remain unchanged (results not shown). Flow crushing gradients were also not used here, which have been shown to increase the measured ATTs across the brain (Dai et al., 2017), though the spoiler gradients sandwiching the GRASE refocussing pulses will have caused some flow crushing (Günther et al., 2005). Since resting ASL scans do not typically use a visual stimulus and vascular crushing is not currently recommended for clinical scans (Alsop et al., 2015), it is likely an ATT prior range of 0.5 - 1.8 s is sufficient for protocol optimisation for young healthy volunteers. However, a range of 0.5 - 2 s may be more appropriate if vascular crushing is used or for older populations (Dai et al., 2017).

### 6.3. Variable noise across ATT

There was a gradual increase in the in vivo CBF posterior distribution SDs at shorter ATTs compared to the simulations, which assumed equal noise across all ATTs. One explanation is that shorter ATTs are generally located closer to the middle of the brain and so further from the head-coil receive elements than longer ATTs. This could result in an SNR level that was negatively corelated with ATT. Another explanation is that shorter ATTs are in regions closer to larger upstream arteries and so experience greater signal variability due to cardiac pulsation.

To test the hypothesis that the CBF posterior SDs vary with ATT, the voxelwise temporal noise, σ, was calculated from the calibrated in vivo single-PLD control images by taking the SD across repeats. A linear model, σ(ATT) = *a* · ATT + *b*, was fit to these data from all subjects using the ground-truth ATT estimates and the “fit” function in MATLAB using bisquare weights, which is robust to outliers. The fitted parameters were *a* = −4.28 × 10^−4^ s^−1^ and *b* = 20.29 × 10^−4^ with the model explaining 58% of the variance (*R*^2^ = 0.58), indicating there is increased noise in the control images at locations with shorter ATTs.

This noise model was used in additional MC simulations, similar to those described in the Methods section, after being rescaled so that σ(1.25 s) was equal to the noise SD used in the original simulations. Figure 8 shows a comparison between the variable noise simulations and in vivo data, demonstrating a much-improved qualitative match than the fixed noise simulations. This suggests the relationship between ATT and temporal signal variation largely explains the differences seen in the trends in vivo, though we cannot deduce the cause of this variability. This ATT dependent noise model is unlikely to be useful for protocol optimisation because it will likely vary across subjects, subject placement, and head coil design.

**Figure 8:**
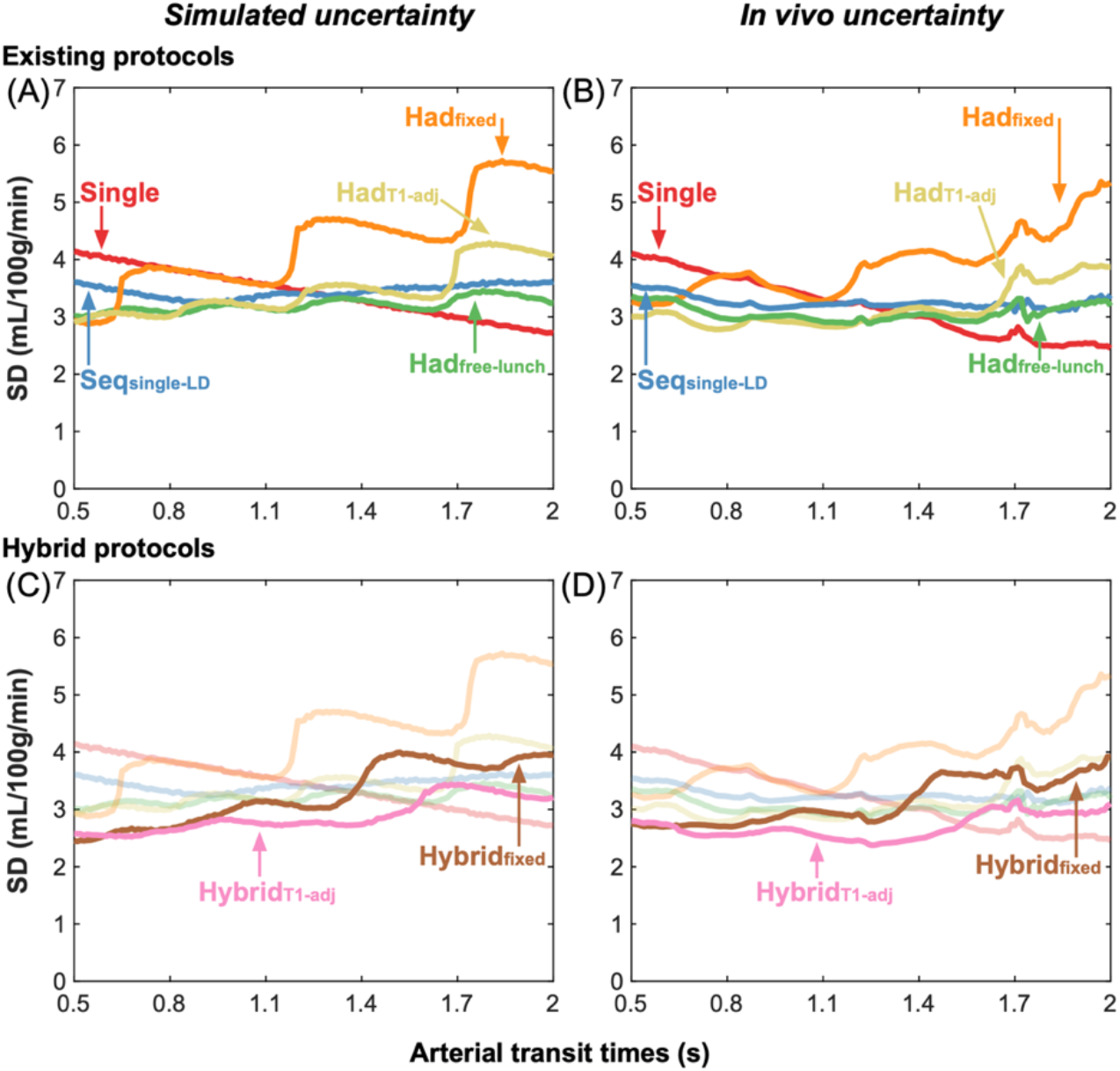
The simulation (Monte Carlo simulation posterior SDs) (A, C) and in vivo (posterior SDs) (B, D) median CBF uncertainty measures for the literature protocols (top) and the proposed hybrid protocols (bottom). These MC simulations use the estimated variable noise levels across ATTs calculated from the in vivo data and scaled to the noise SD used in the original MC simulations. The simulation uncertainty trends across ATTs now better match the trends seen in vivo.

### 6.4. Subject CBF and ATT variation

Large differences in the CBF and ATT maps were seen across subjects (Supporting Information Figure S2 and Supporting Information Figure S3). These differences may be due to previously seen global variations across age and sex, such as decreasing CBF and increasing ATT with age (Chen et al., 2011; Dai et al., 2017; Parkes et al., 2004) and higher CBF and lower ATT in women (Henriksen et al., 2013; MacIntosh et al., 2010; Vernooij et al., 2008).

### 6.5. Long label duration protocols

The longest LD used in this work was 1.8 s, which is currently recommended for clinical use with single-PLD (Alsop et al., 2015). It has been suggested that it is more SNR efficient to use long LDs of 3 - 4 s (resulting in fewer averages) for single-PLD PCASL (Zun et al., 2014) with additional benefits of reduced temporal signal variation and reduced sensitivity to delayed ATTs (Dai et al., 2012; Lebel et al., 2015). To investigate the extent to which the CBF accuracy of the protocols in this work may benefit from longer LDs, we repeated the protocol optimisations with a maximum LD of 5 s and conducted further MC simulations as before. The LD of the single-PLD protocols was optimised similar to (Zun et al., 2014) but for a 5 minute scan, a PLD of 2 s, and an ATT range 0.5 - 2 s.

The optimised protocol timings are given in Supporting Information Table S2 and the MC simulation fitting posterior distribution SDs are shown in Figure 9. All the protocols used much longer LDs than when the maximum LD was 1.8 s, except Had*_T1_*_-adj_ which had the same timings as before. The increase in the protocols’ LDs led to an average reduction in the CBF posterior SDs of 0.17 ± 0.07 mL/100g/min (5.1% ± 1.9%). As before, the Hybrid*_T1_*_-adj_ and Hybrid_variable_ protocols had similar posterior SDs, which were the lowest of all the protocols, including single-PLD. It is possible that the in vivo benefits of using longer LDs extend beyond the theoretical benefits found here (Dai et al., 2012; Lebel et al., 2015) and should be investigated further.

**Figure 9:**
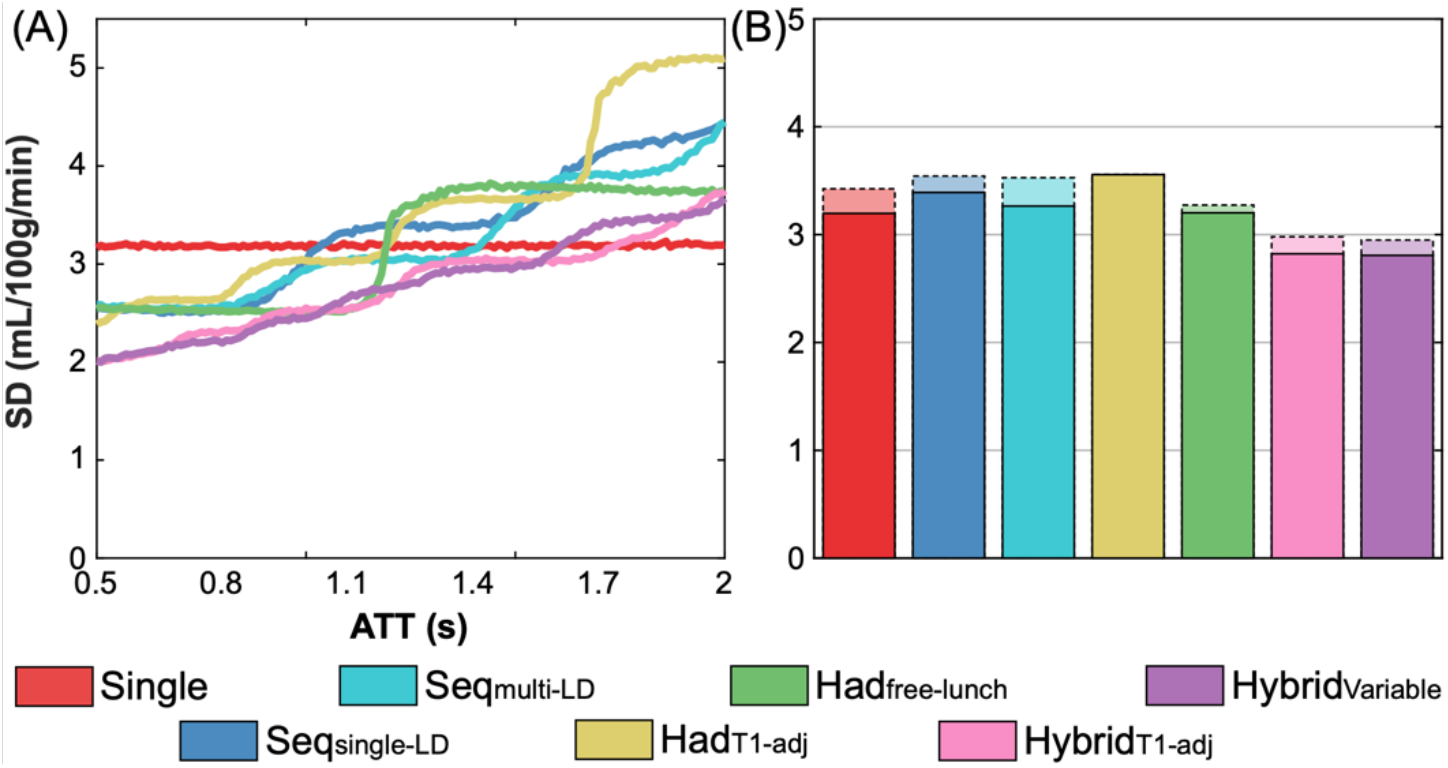
The MC simulation CBF posterior SDs (uncertainty) for a selection of the protocols optimized using longer LDs. (A) the median uncertainty for each protocol across ATTs, (B) the mean uncertainty for each protocol across the whole ATT range. The mean uncertainties for the short LD cases are also shown in (B) as faded bar graphs to demonstrate the differences.

## 7. Conclusions

In this work, we demonstrated that optimised multi-timepoint protocols can generate more confident, accurate, and repeatable CBF estimates across a given ATT range than a single-PLD protocol, while also generating ATT maps. We found that the time-encoded free-lunch protocol with *T_1_*-adjusted LDs can lead to improved CBF estimates over a fixed-LD time-encoded protocol and is a good approximation to the optimal time-encoded design. Finally, we demonstrated that a novel hybrid time-encoded with sequential PLD protocol design utilising *T_1_*-adjusted label durations out-performed a wide range existing literature protocol designs for estimating CBF, both in simulation and in vivo.

## 10. Supporting information text 1 - Protocol optimisation

### Optimisation updates

Details of the optimisation implementations are as follows. To optimise the Seq_single-LD_ protocol, the PLDs were optimised in the same way as the original implementation, but the optimal single experiment LD was found when optimising the final (*N*^th^) PLD. For the Seq_multi-LD_ protocol, the optimal i^th^ LD and PLD pair were found at each iteration. In both cases, the PLDs were restricted to a monotonically increasing order (PLD_i-1_ ≤ PLD_i_ ≤ PLD_i+1_) to reduce the parameter space.

Due to the design constraints of the Had_fixed_ and Had*_T1_*_-adj_ protocols, only the first LD and final PLD, for any encoding size, (*M* + 1) × *M*, must be searched over, making it possible to carry out a global grid search for all possible timing combinations for each *M*, rather than use the iterative exchange method used with the sequential protocols. The Had_free-lunch_ protocol differs only in that the first encoded LD is fixed to the single-PLD protocol LD, with the remaining LDs being optimised identically to the Had_fixed_ and Had*_T1_*_-adj_ protocols.

The Hybrid_fixed_ and Hybrid*_T1_*_-adj_ protocols were optimised by iterating through each of the sequential *N* PLDs and optimising the i^th^ PLD and LDs of the encoding matrix simultaneously.

Had_variable_ was optimised by iterating through the encoded LDs and simultaneously optimising the i^th^ LD and the final PLD. Hybrid_variable_ was optimised by iterating through each of the sequential *N* PLDs and encoding matrices and then iterating through each of the *M* LDs, optimising the j^th^ LD of the i^th^ encoding matrix with the i^th^ PLD together. Initial testing of these variable-LD protocols suggested the best protocols had LDs of decreasing duration during the PCASL preparation, so the LD was restricted to LD_j-1_ ≤ LD_j_ ≤ LD_j+1_.

### Protocol initialisations

Seq_single-LD_: initialised with single LD 1.8 s and *N* PLDs spaced evenly between 0.075 - 2.3 s. Seq_multi-LD_: *N* LD and PLD pairs randomly initialised between 0.8 - 1.8 s and 0.075 - 2.3 s, respectively. For each *N*, the sequential protocol optimisations iterated through each timepoint in a randomly permuted manner and were run with 20 different initialisations for robustness.

Had_free-lunch_: the first LD was fixed at 1.8 s (matching the single-PLD protocol) with the remaining LDs being either fixed-duration or *T_1_*-adjusted. The final PLD was also optimised, therefore, it was not guaranteed that the PLD of the first encoded LD would match that of the single-PLD protocol. The Had_fixed_ and Had*_T1_*_-adj_ protocols did not require initialisation because the entire timing parameter space could be evaluated.

Hybrid_fixed_ and Hybrid*_T1_*_-adj_: all *N* final PLDs initialised at 0.075 s - the LDs did not require initialisation because they are all globally optimised at each step, similar to the time-encoded protocols.

Had_variable_ and Hybrid_variable_: the LDs were randomly initialised between 0.1 - 1.8 s and sorted into a descending order; the *N* PLDs were initialised to 0.075 s. In the case of Hybrid_variable_, the *N* PLDs were iterated through in the same order. For both Had_variable_ and Hybrid_variable_, the *M* LDs were iterated through in a randomly permuted order. The optimisations were each run with 50 different initialisations for robustness.

## 11. Supporting information Figures and Tables

**Supporting Information Figure S1:**
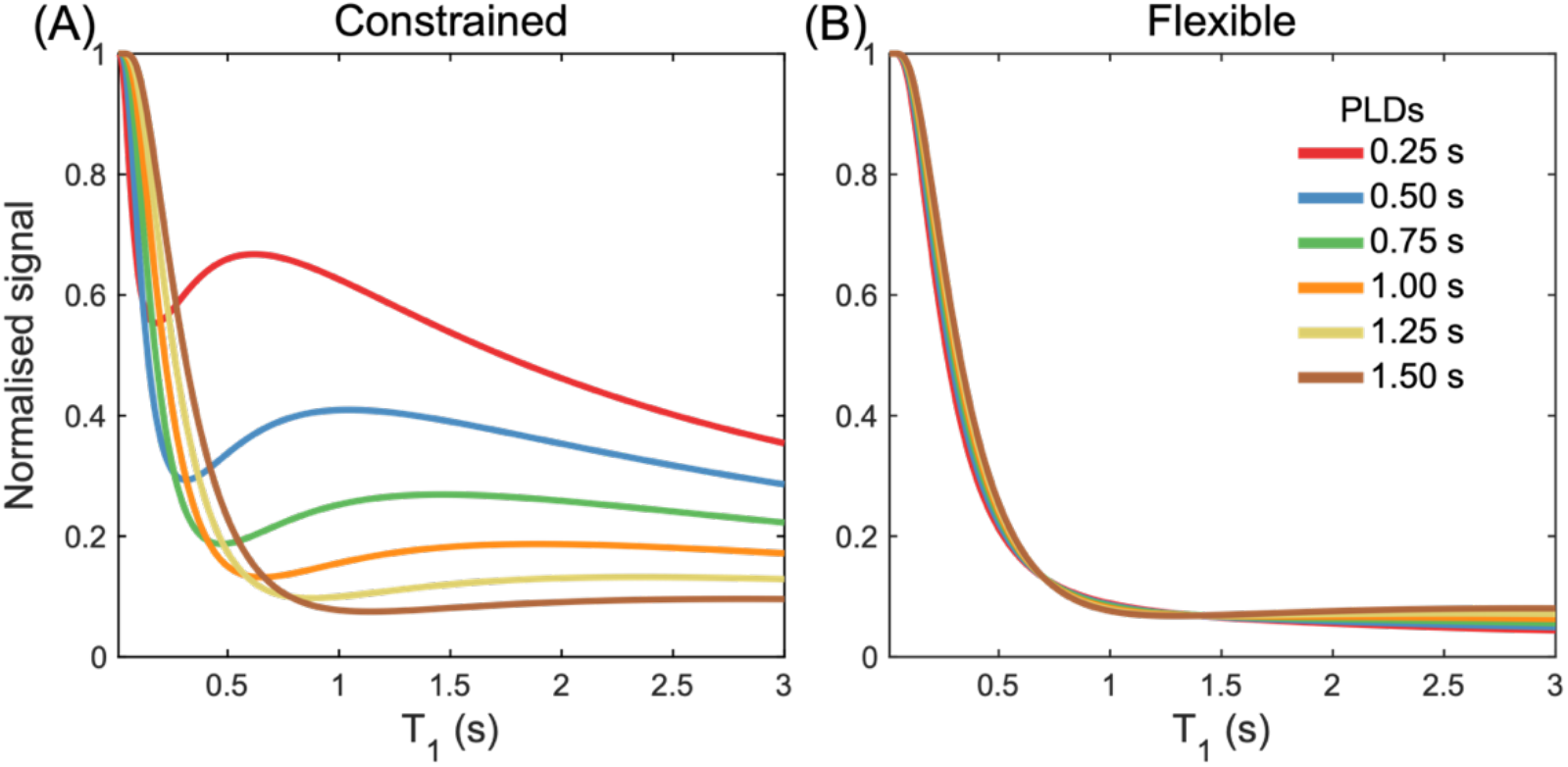
The theoretical residual static tissue longitudinal magnetisation at the time of the readout excitation. The BGS uses a presaturation module and two inversion pulses to. null T_1_ = 700 ms and 1400 ms. (A) The residual magnetisation when the inversion pulses are restricted to play out after the LD; (B) the residual magnetisation when the inversion pulses are played out at the optimal times, including during the LD. For both (A) and (B), the residual longitudinal magnetisation is shown for 6 different PLDs (0.25 - 1.5 s) and a LD of 1.4 s. Instantaneous RF pulses, perfect spoiling, and perfect inversion are assumed. The null time has been set to 100 ms before the excitation, to ensure positive signal in all cases.

**Supporting Information Figure S2:**
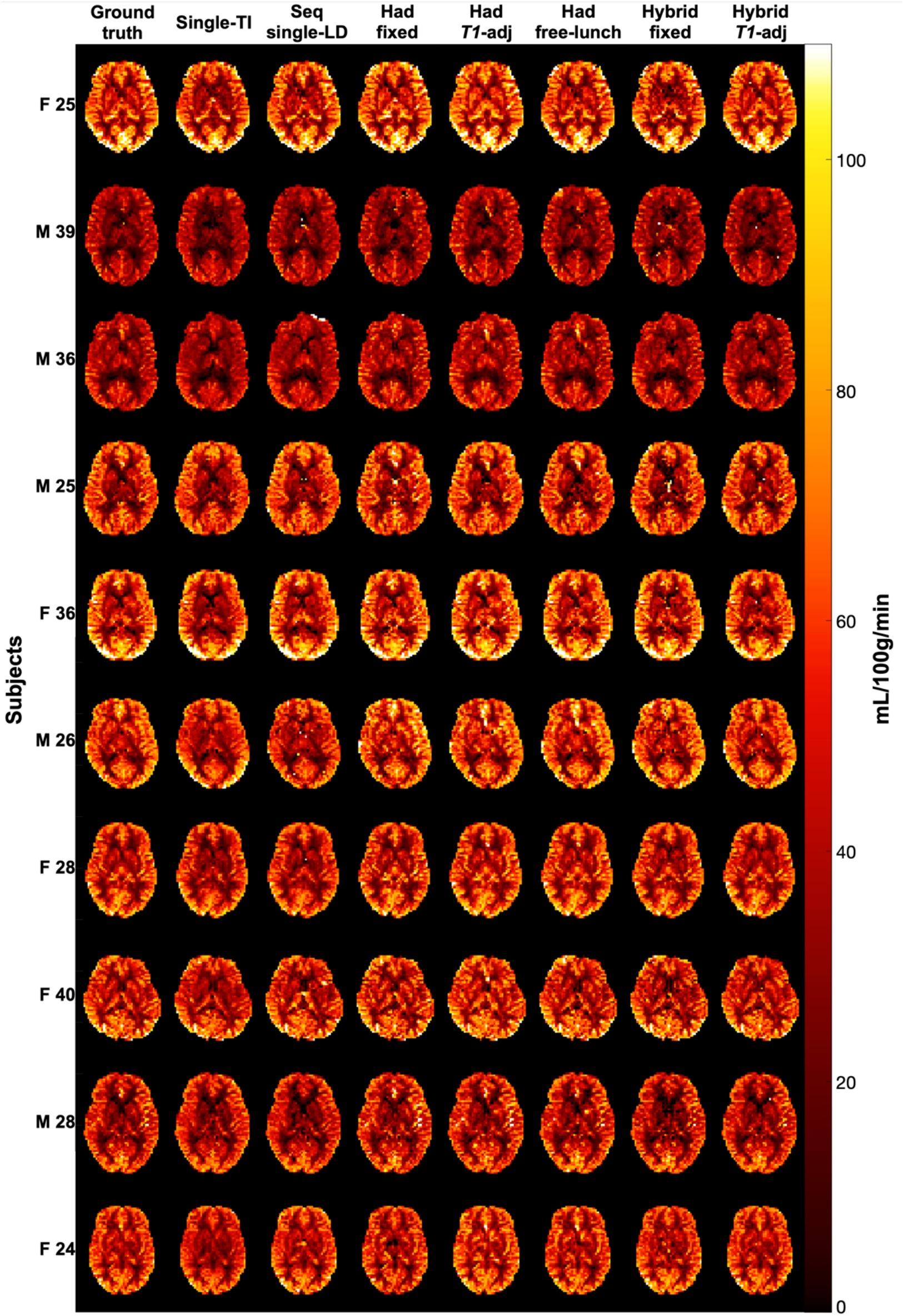
A single slice of the CBF maps for each subject for each of the protocols and the ground truth estimates. The subjects’ sex and age are given, where “F 25” means “Female, 25 years old.”

**Supporting Information Figure S3:**
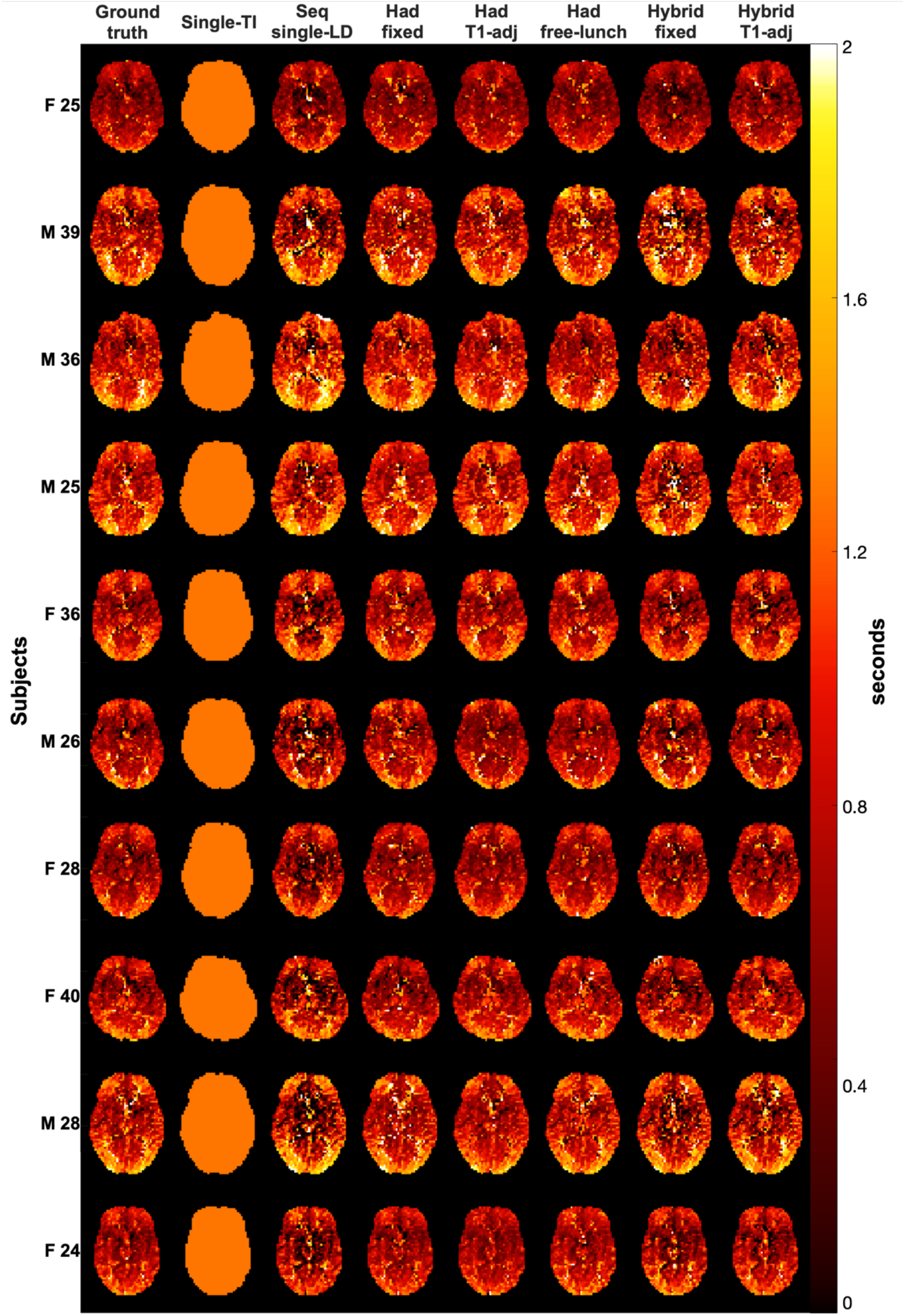
A single slice of the ATT maps for each subject for each of the protocols and the ground truth estimates. The subjects’ sex and age are given, where “F 25” means “Female, 25 years old.”

**Supporting Information Figure S4:**
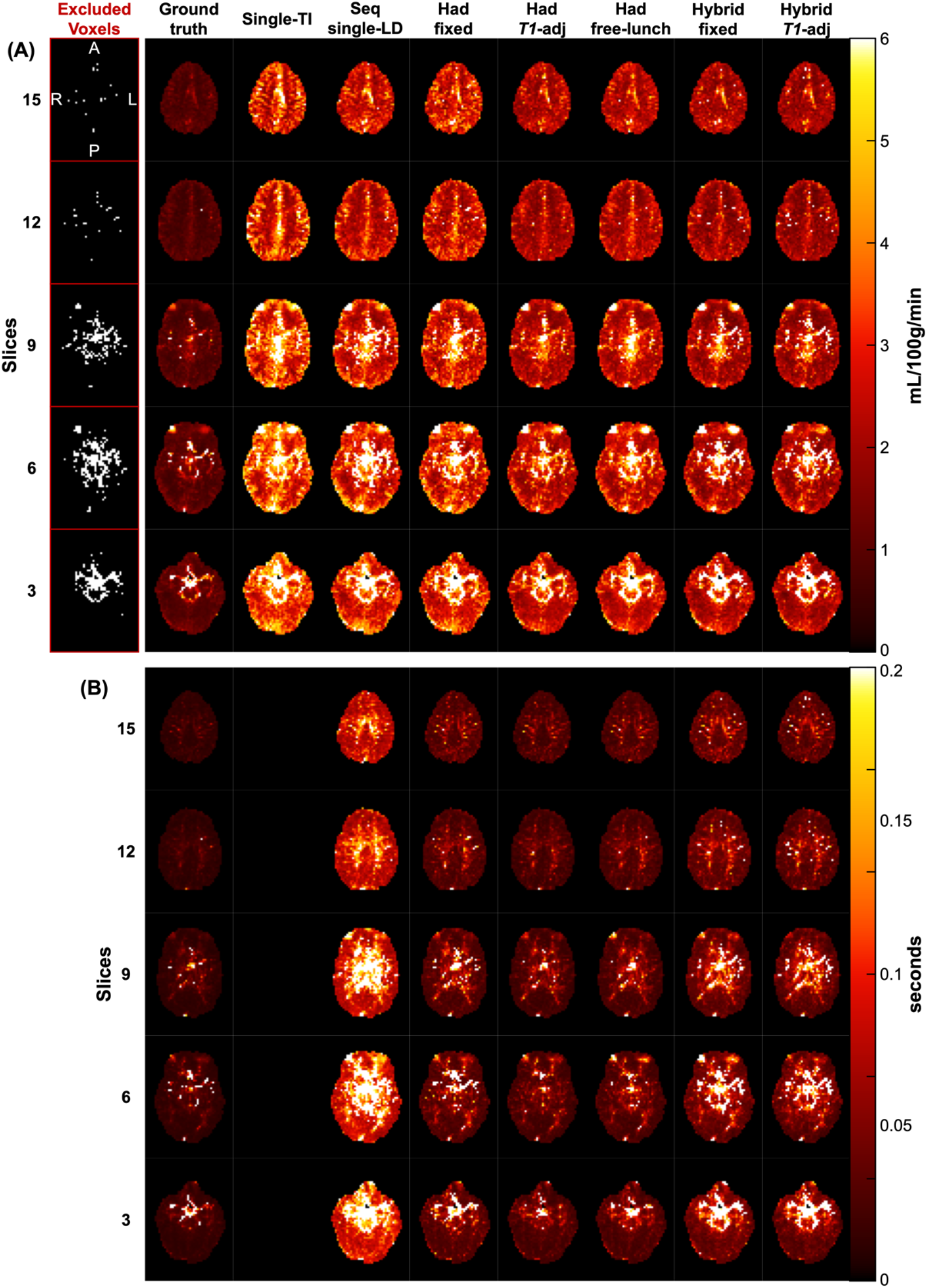
The voxels excluded due to the posterior SD restrictions and the posterior SD maps for 5 slices of a single representative subject. (A) The excluded voxels and the CBF posterior SD maps, (B) the ATT posterior SD maps. The single-PLD protocol does not have ATT posterior SD maps because ATT is not estimated. The excluded voxel maps show voxels excluded due to high SDs in either the CBF or the ATT posterior SD maps.

**Supporting Information Figure S5:**
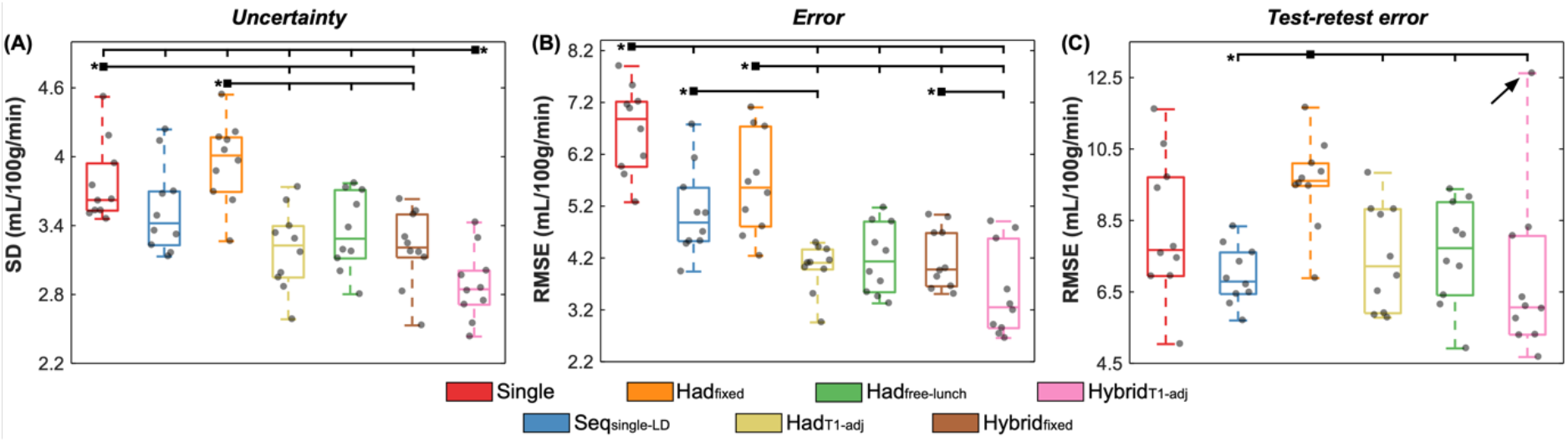
The subjectwise in vivo comparisons of the uncertainty (posterior SDs), accuracy (RMSEs), and repeatability (test-retest RMSEs) metrics for each protocol. The boxplots show the median, quartiles, and range across subjects, in each case. Significant differences are shown for individual protocols (two-sided paired Wilcoxon signedrank test, Bonferroni correction for 6 comparisons, α<0.05).

**Supporting Information Figure S6:**
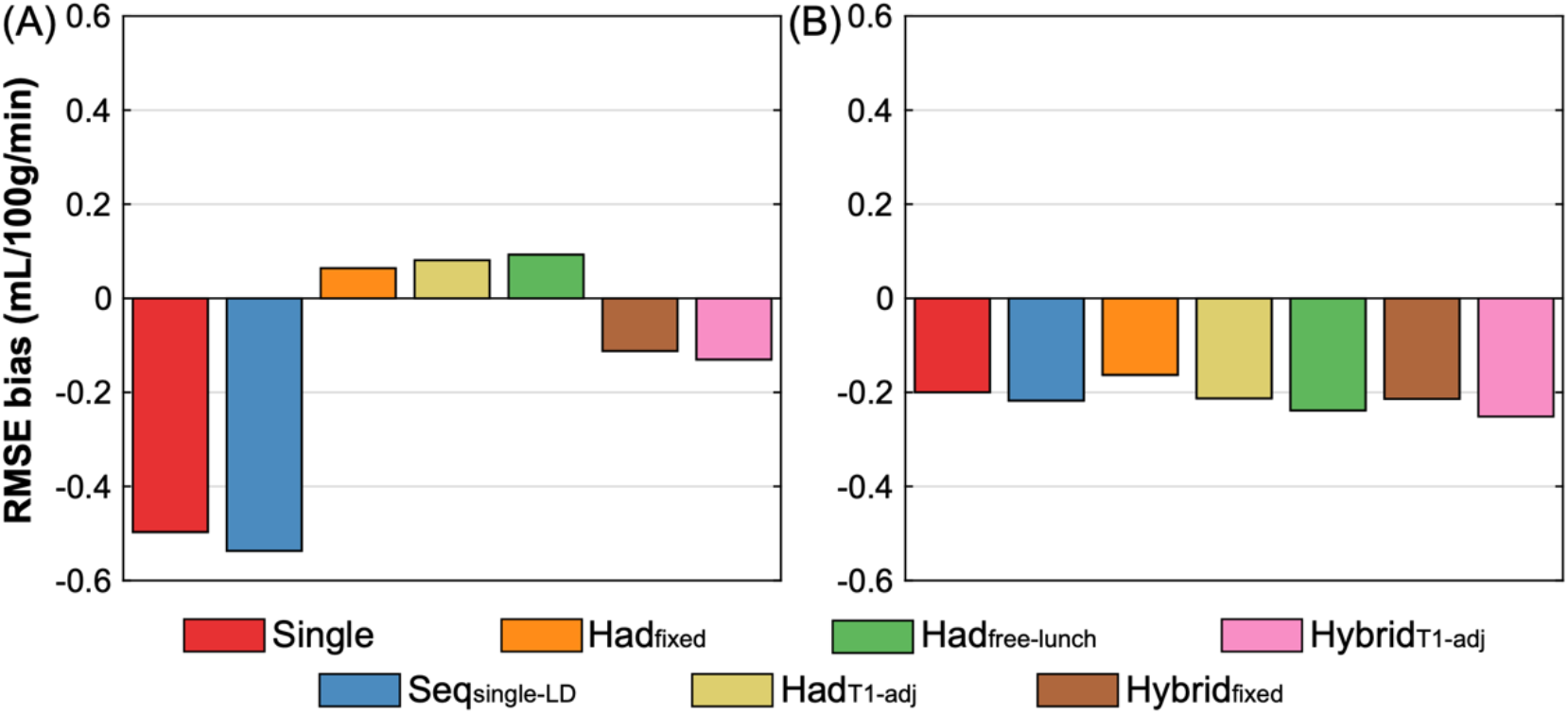
Bias in the ground truth Monte Carlo simulation CBF RMSEs when using 2 different noise models: (A) ground truth values fitted using 1 noise magnitude for all of the data and (B) ground truth values fitted using 3 noise magnitudes (1 each for: non-time-encoded protocols, time-encoded protocols, and the hybrid protocols). When 1 noise magnitude is used in the fitting, there is a large variation in the bias across protocols, but when 3 noise magnitude are used the RMSEs are much more similarly underestimated for all the protocols by -0.21 ± 0.03 mL/100g/min.

**Supporting Information Figure S7:**
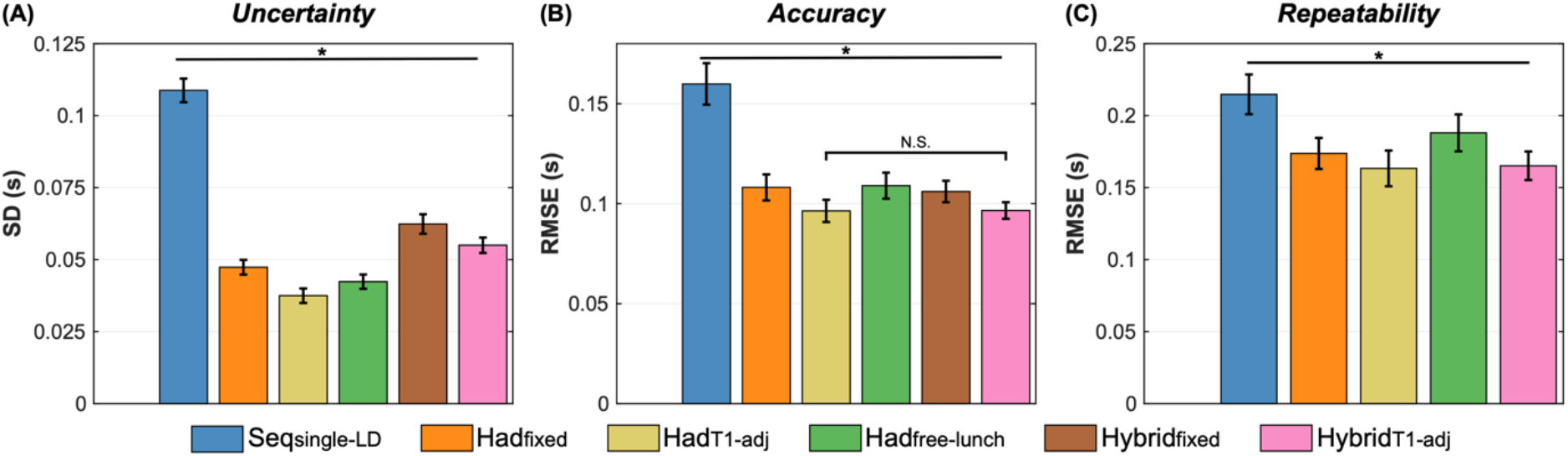
The in vivo voxelwise ATT measures of (A) uncertainty (posterior distribution SDs), (B) accuracy (RMSEs relative to the ground truth estimates) and (C) repeatability (test-retest RMSEs). The mean and standard error (see methods) of the metrics across voxels are shown. All protocols had significantly different measures, unless highlighted as not-significant (NS).

**Supporting Information Figure S8:**
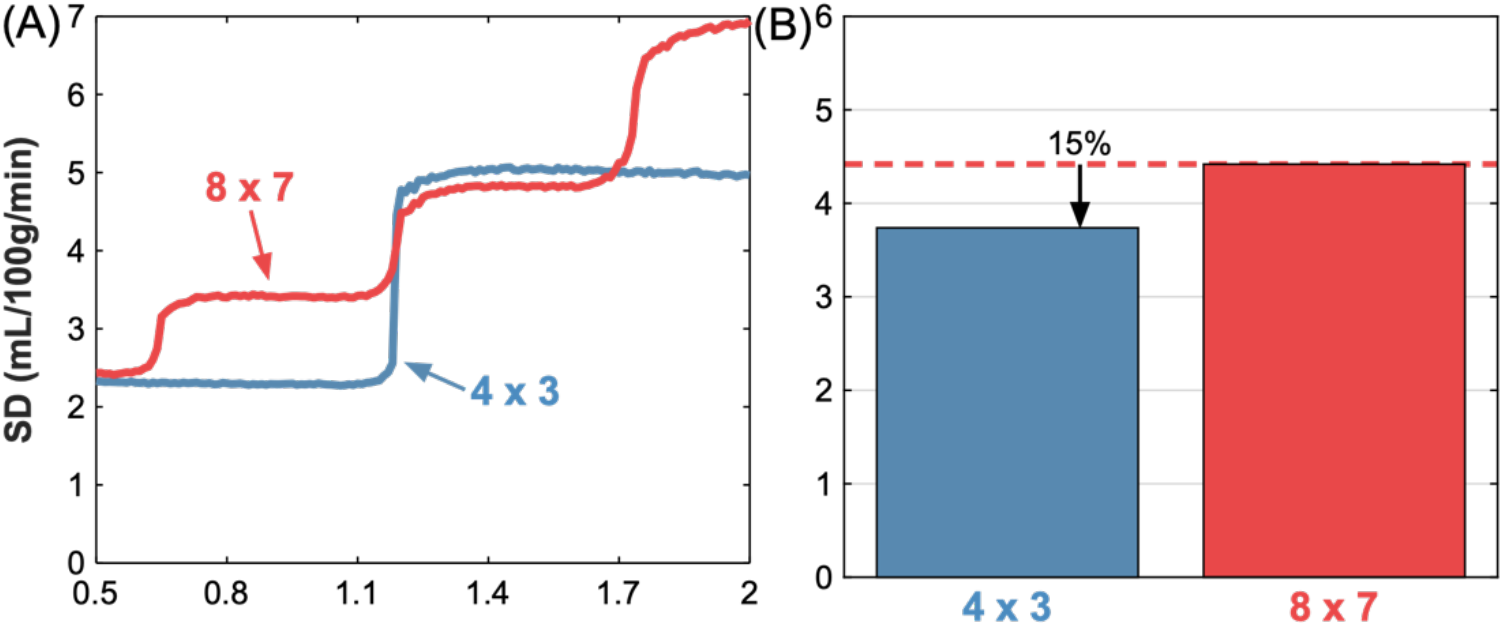
The simulated CBF uncertainty (Monte Carlo simulation posterior SDs) for the optimised Had_fixed_ protocol when using a 4×3 or 8×7 Hadamard matrix. (A) the median uncertainty across ATTs, (B) the mean uncertainty for each encoding size across the whole ATT range. Due to the 4×3 Hadamard protocol only having 3 PLDs, there is a large increase in uncertainty in the middle of the ATT distribution where the protocol becomes more sensitive to longer ATTs. However, overall, the 4×3 protocol has a lower average uncertainty than the 8×7 protocol.

**Supporting Information Figure S9:**
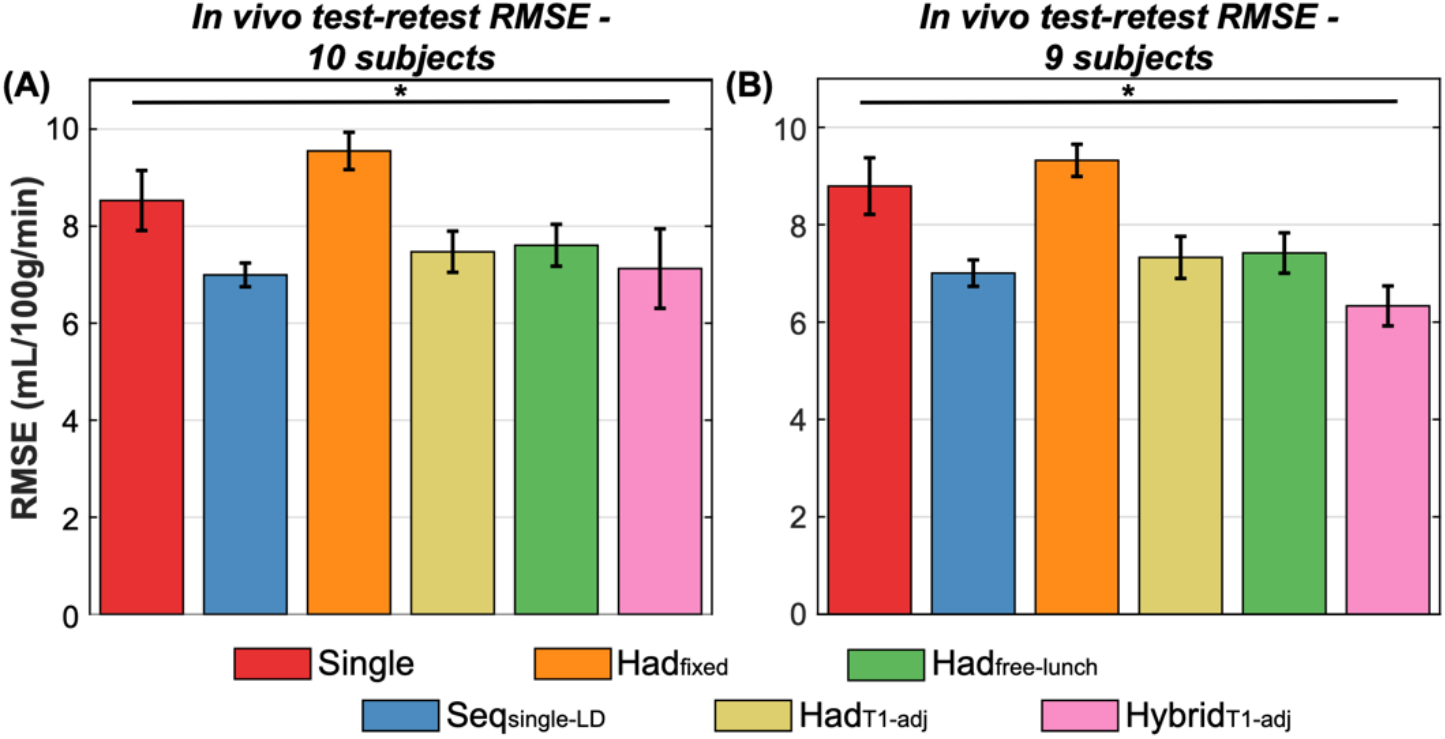
The in vivo test-retest RMSEs across all voxels with all subjects (A) and with 1 subject removed (B). The removed subject had a much larger Hybrid_T1-adj_ test-retest RMSE than the other subjects, but when removed did not lead to a large change in the test-retest RMSEs of the other protocols. The means and standard errors of the bootstrap distributions are shown (see methods). All differences were significant (two-sided paired Wilcoxon signed-rank test, Bonferroni correction for 6 comparisons, α<0.05).

**Supporting Information Table S1:** The optimised protocol timings for Seq_multi-LD_ and Had_fixed_ with a 4×3 Hadamard matrix, which were not included in the in vivo comparison. N_T_ is the number of effective PLDs, N_Ave_ is the number of averages, and N_acq_ is the number of acquired volumes for each scan.

| Protocol | Label durations (ms) | Post-label delays (ms) | $N_T$ | $N_{Ave}$ | $N_{Acq}$ | Scan duration (min) |
| --- | --- | --- | --- | --- | --- | --- |
| *Seq<sub>multi-LD</sub>* | 1800, 1500, 1800, 1800, | 200, 825, 1350, 1475, 1800, | 12 | 3 | 72 | 5:00 |
|  | 1800, 1800, 1800, 1800, | 1850, 2100, 2225, 2300, |  |  |  |  |
|  | 1800, 1800, 1800, 1800 | 2300, 2300, 2300 |  |  |  |  |
| *Had<sub>fixed</sub>*<br>*4×3* | 1125, 1125, 1125 | 75 | 3 | 18 | 72 | 4:54 |

**Supporting Information Table S2:** The optimised protocol timings when the maximum LD was extended to 5 s. For the time-encoded (Had) and hybrid protocols, the LDs are given in chronological order and the number of LDs defines the size of the Hadamard encoding matrix used. For the Hybrid_variable_ protocol, each PLD is associated with the LDs on the same row. N_T_ is the number of effective PLDs, N_Ave_ is the number of averages, and N_acq_ is the number of acquired volumes for each scan.

| Protocol | Label durations (ms) | Post-label delays (ms) | $N_T$ | $N_{Ave}$ | $N_{Acq}$ | Scan duration (min) |
| --- | --- | --- | --- | --- | --- | --- |
| Single-PLD | 3125 | 2000 | 1 | 26 | 52 | 5:00 |
| Seqsingle-LD | 2800 | 75, 975, 1600, 2100, 2300, 2300 | 6 | 5 | 60 | 5:00 |
| Seqmulti-LD | 2025, 1400, 3425, 4225, 3850 | 200, 925, 1975, 1475, 2300 | 5 | 6 |  | 5:00 |
| Hadfree-lunch | 3125, 1100, 1100 | 100 | 3 | 12 | 48 | 4:51 |
| Hybrid<sub>T1-adj</sub> | 3550, 1050, 625 | 75, 275, 550, 625 | 12 | 3 | 48 | 5:00 |
| Hybrid<sub>variable</sub> | 3975, 1125, 300<br>3325, 1425, 550<br>3300, 1225, 500<br>3575, 725, 625 | 200<br>325<br>575<br>675 | 12 | 3 | 48 | 5:00 |

